# Multi-omic analyses identify molecular targets of Chd7 that mediate CHARGE syndrome model phenotypes

**DOI:** 10.1101/2025.07.28.666396

**Authors:** Melody B Hancock, Dana R Ruby, Rachael A Bieler, David C Cole, Kurt C Marsden

## Abstract

CHARGE syndrome is a developmental disorder that affects 1 in 10,000 births, and patients exhibit both physical and behavioral characteristics. *De novo* mutations in *CHD7 (chromodomain helicase DNA binding protein 7)* cause 67% of CHARGE syndrome cases. CHD7 is a DNA-binding chromatin remodeler with thousands of predicted binding sites in the genome, making it challenging to define molecular pathways linking loss of *CHD7* to CHARGE phenotypes. To address this problem, here we used a previously characterized zebrafish CHARGE model to generate transcriptomic and proteomic datasets from larval zebrafish head tissue at two developmental time points. By integrating these datasets with differential expression, pathway, and upstream regulator analyses, we identified multiple consistently dysregulated pathways and defined a set of candidate genes that link loss of *chd7* with disease-related phenotypes. Finally, to functionally validate the roles of these genes, CRISPR/Cas9-mediated knockdown of *capgb*, *nefla*, or *rdh5* phenocopies behavioral defects seen in *chd7* mutants. Our data provide a resource for further investigation of molecular mediators of CHD7 and a template to reveal functionally relevant therapeutic targets to alleviate specific aspects of CHARGE syndrome.

**Summary Statement:** We have identified Chd7 target genes *capgb*, *nefla*, and *rdh5* that mediate CHARGE model phenotypes from transcriptomic and proteomic analysis of *chd7* wild type, heterozygous, and homozygous mutant zebrafish brain tissue at two developmental time points.

## Introduction

CHARGE syndrome is an autosomal dominant developmental condition that occurs 1 in every 10,000 newborns (Blake et al., 1998) and is named for its previous diagnostic criteria: C-coloboma, H-heart disease, A-atresia choanae, R-retardation of growth and development and/or CNS anomalies, G-genital hypoplasia, and E-ear anomalies and/or deafness (Pagon et al., 1981). Affected individuals display physical as well as complex behavioral characteristics such as autistic-like behaviors (Hartshorne et al., 2005), obsessive–compulsive disorder (Hartshorne et al., 2017), attention-deficit/hyperactivity disorder (AD/HD) (Hartshorne and Cypher, 2004), anxiety (Hartshorne et al., 2017), aggression (Blake et al., 2005), intellectual disability (Wachtel et al., 2007), and sensory disorders (Edwards et al., 1995).

67% of CHARGE syndrome cases can be attributed to a mutation in *chromodomain helicase DNA binding protein 7* (*CHD7*) (Zentner et al., 2010). CHD7 is ubiquitously expressed in early human development (Sanlaville et al., 2006), a pattern conserved across species including: mouse (Bosman et al., 2005), zebrafish (Patten et al., 2012), fly (Daubresse et al., 1999), and chick (Williams et al., 2024). CHD7 is part of the CHD family of chromatin remodelers and is recruited to specific histone modifications to enhancer regions to regulate the accessibility of DNA via ATP-dependent chromatin remodeling (Basson and van Ravenswaaij-Arts, 2015). CHD7 ChIP-seq experiments have found 10,000 predicted binding sites in mouse embryonic stem cells (Schnetz et al., 2010) and 9,000 predicted binding sites in granule cell precursors of the mouse cerebellum (Reddy et al., 2021). CHD7 is known to be important in neurodevelopment through its involvement in neural crest cell migration and specification (Schulz, Wehner, et al., 2014), neurogenesis (Feng et al., 2013), neuronal differentiation (Jones et al., 2015), and activation of neural crest transcription factors *sox9*, *twist*, and *slug* (Bajpai et al., 2010). Not only that, but when embryonic stem cells transition to neural progenitors CHD7 binding patterns change, suggesting that CHD7’s functions shift across neurodevelopmental stages (Schnetz et al., 2009).

CHARGE models have been established in systems including mouse, fly, and zebrafish, and these have been studied to determine how loss of *chd7* affects multiple aspects of development: brain defects (Whittaker, Riegman, et al., 2017), (Feng et al., 2017), (Donovan et al., 2017), ear defects (Bosman et al., 2005) (Gao et al., 2024), eye defects (Gage et al., 2015), cardiac defects (Bergman et al., 2010) (Bosman et al., 2005), and craniofacial defects (Asad et al., 2016), (Asad and Sachidanandan, 2020), (Balow et al., 2013). These models have also shown how loss of *chd7* affects sensory systems (Bergman et al., 2010) (Layman et al., 2009) (Bosman et al., 2005), behavior (Hodorovich et al., 2023) (Jamadagni et al., 2021), and the transcriptome (Jamadagni et al., 2021), (Yao et al., 2020), (Stathopoulou et al., 2023), (Huang et al., 2025). Despite these advances in understanding the biological roles of CHD7, the molecular mechanisms that directly link loss of *chd7* with CHARGE syndrome phenotypes remain unclear.

In this study we aimed to leverage the advantages of our established zebrafish CHARGE model (Hodorovich et al., 2023) to investigate the molecular pathophysiology of neurobehavioral symptoms associated with CHARGE syndrome. By integrating transcriptomic and proteomic analyses across two developmental timepoints, we identify key *chd7*-dependent genes, proteins, and pathways, including histone modification, GABAergic receptor, NMDA receptor, ROBO receptor, and SNARE signaling. Finally, we filtered our datasets to identify individual candidate mediators that were consistently dysregulated at the RNA and protein level, at both timepoints, and in both *chd7* heterozygotes and homozygous mutants. We then cross-referenced these with known CHD7 binding targets, and using CRISPR/Cas9-mediated knockdown, we functionally validated *capgb*, *nefla*, and *rdh5* as likely mediators CHARGE model behavioral and morphological phenotypes.

## Results

To identify molecular mediators of Chd7-dependent development, we collected head tissue at 3- and 5-days post fertilization (dpf) from zebrafish larvae derived from incrosses of *chd7* heterozygotes. These timepoints represent the earliest developmental stages when CHARGE-related morphological and behavioral phenotypes emerge (Hodorovich et al., 2023). By analyzing both timepoints we also sought to gain insight into how Chd7 modulates developmental gene expression patterns across these key stages. Tail tissue was used for genotyping, and head tissue from larvae (n=20-30 per genotype) was then pooled for subsequent RNA or protein isolation to generate 4 biological replicates for all three genotypes: *chd7* wildtype (WT), heterozygous (HT), and mutant (MUT) samples. Following RNA sequencing and mass spectrometry using standard methods as outlined in the Methods section, we performed differential expression analysis comparing HT to WT samples (HT v WT) and MUT to WT samples (MUT v WT) to identify altered gene and protein levels due to loss of *chd7*. First, to determine the degree of similarity in overall RNA and protein expression between replicates, timepoints, and genotypes, we used principal component analysis (PCA). PCA of the transcriptome data shows that samples from all genotypes cluster by timepoint, and PC1 alone accounts for 82% of the variance between samples (Figure 1A). PCA of the proteome data reveals a similar clustering by timepoint, with PC1 accounting for 34.4% and PC2 23.8% of the variance (Figure 1B). While RNA samples show little clustering by *chd7* genotype, protein samples do cluster somewhat by genotype, particularly at 5 dpf. Overall, developmental changes in gene expression from 3 to 5 dpf are the primary driver of the differences between samples, but by 5 dpf Chd7-dependent molecular changes can be seen by PCA.

**Figure 1.**
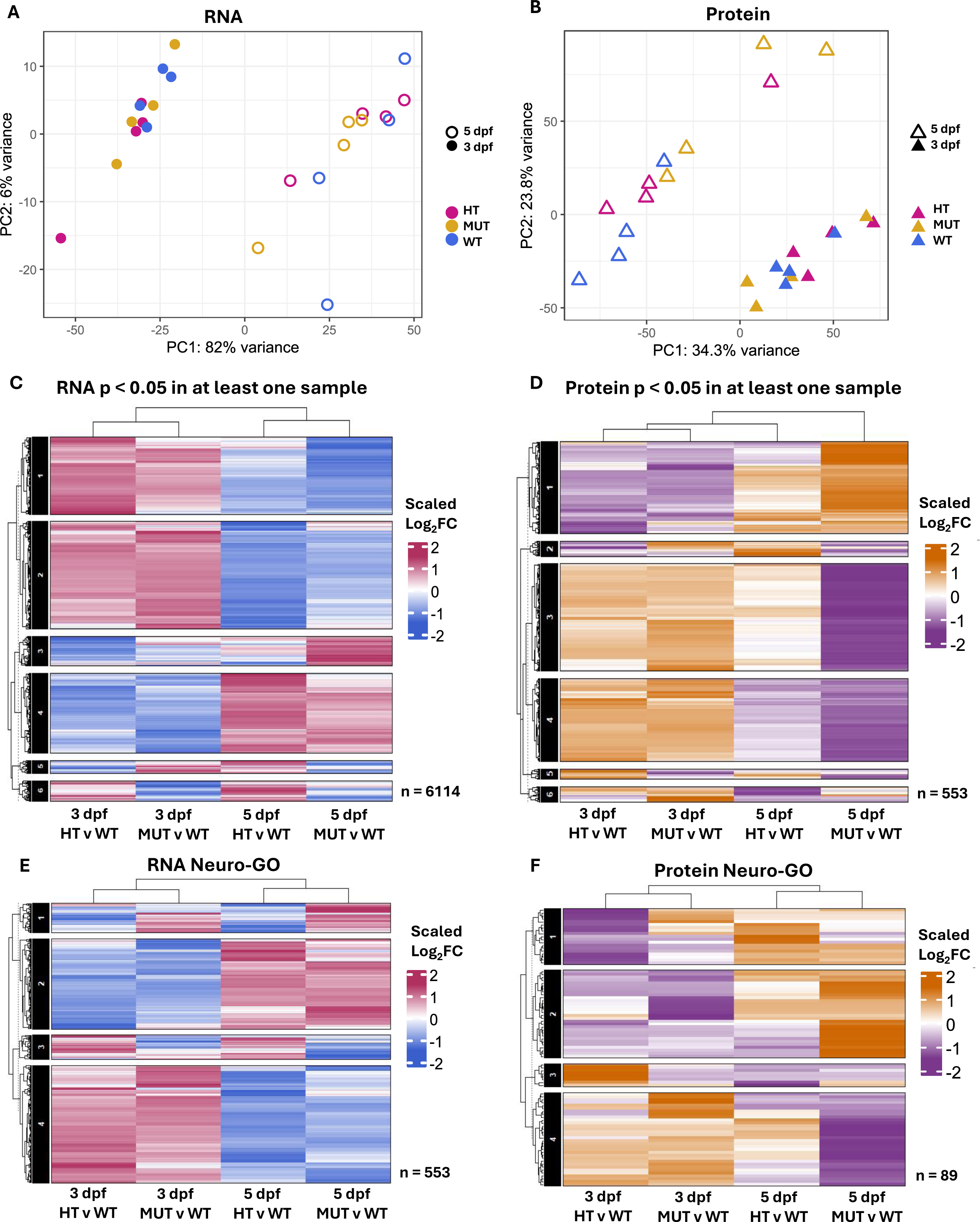
Loss of *chd7* causes dysregulation of genes associated with neurodevelopmental and transcriptional regulation functions. Principal component analysis (PCA) of **A**. quantified RNAs for each sample and replicate, and **B**. quantified proteins for each sample and replicate. Heatmap of scaled Log_2_ Fold Change (Log_2_FC) with hierarchical clustering by similar fold change patterns of **C**. Differentially Expressed Genes (DEGs) with a p-value < 0.05 in at least one sample (n = 6114) and **D**. Differentially Expressed Proteins (DEPs) with a p-value < 0.05 in at least one sample (n = 553). Heatmap of scaled Log_2_FC with hierarchical clustering by similar fold change patterns of **E**. DEGs that overlap with a list of neurodevelopmental genes generated from GO Accession terms and Ensembl (Neuro-GO) (n = 553) and **F**. DEPs that overlap with the Neuro-GO list (n=89). **C-F**. Slices (left, numbered in black and white) by hierarchical clustering of similar fold change patterns.

Next, we used hierarchical clustering to find genes with similar expression patterns across timepoints and genotypes. We first identified significantly differentially expressed RNAs (DEGs) and proteins (DEPs) in HT and MUT samples compared to WT, and we then generated heatmaps to visualize these differentially expressed genes. We plotted RNAs (Figure 1C) and proteins (Figure 1D) significant (p-value less than 0.05) in at least one sample and performed k-means hierarchical clustering by Euclidean distance (Gu, 2022). DEGs again clustered first by timepoint, with similar overall patterns in HT and MUT. DEPs, however, showed slightly different overall clustering, with 5 dpf HT expression showing more similarity to 3 dpf samples than 5 dpf MUT.

DEGs and DEPs were also clustered by similarity in Fold Change (FC), with DEGs in slices 1, 2, and 4 showing similar patterns by timepoint and those in slice 6 having similarity by genotype, being largely upregulated in both 3 and 5 dpf HT and downregulated in 3 and 5 dpf MUT (Figure 1C). Slice 6 includes 381 unique RNAs significantly (FDR) associated with GO biological processes: chromosome condensation (GO:0030261), negative regulation of DNA recombination (GO:0045910), and chromatin remodeling (GO:0006338) (Supplemental Table 1). DEP clustering revealed a set of 18 proteins in slice 5 with similarity by *chd7* genotype, mostly upregulated in HT and downregulated in MUT at both timepoints. Slice 5 includes 18 unique proteins not significantly (FDR) associated with GO biological processes: hydrogen sulfide metabolic process (GO:0070813), carnitine metabolic process CoA-linked (GO:0019254), and positive regulation of ERAD pathway (GO:1904294) (Supplemental Table 2).

To then identify CHD7-mediated patterns in neurodevelopmental gene expression, we generated a list of neuro-related genes from Gene Ontology (GO) Accession terms and Ensembl (Supplemental Table 3). Overall, these neuro-related DEGs and DEPs clustered by timepoint (Figure 1E, F) and based on FC similarity we generated 4 slices with similar patterns. Neuro-related DEGs in slices 1 and 3 show similarity by genotype, with slice 1 displaying upregulation in HT and downregulation in MUT and slice 3 having the opposite pattern (Figure 1E). Slice 1 includes 62 unique RNAs significantly (FDR) associated with GO biological processes: visual perception (GO:0007601), sensory perception of light stimulus (GO:0050953), regulation of G protein-coupled receptor signaling pathway (GO:0008277), and phototransduction (GO:0007602) (Supplemental Table 4). Slice 3 includes 52 unique RNAs significantly (FDR) associated with GO biological processes: positive regulation of synaptic transmission glutamatergic (GO:0051968), neurotransmitter receptor diffusion trapping (GO:0099628), and postsynaptic neurotransmitter receptor diffusion trapping (GO:0098970) (Supplemental Table 5). For neuro-related DEPs, there were no consistent patterns by genotype that distinguish HT from MUT, but proteins in slice 3 were largely downregulated in both 3 and 5 dpf MUT (Figure 1F). Slice 3 includes 8 unique proteins significantly (FDR) associated with GO biological processes: nervous system development (GO:0007399), system development (GO:0048731), multicellular organism development (GO:0007275) (Supplemental Table 6).

In summary, we find that loss of either 1 or 2 copies of *chd7* produces globally similar changes in RNA and protein expression patterns, and these patterns largely reverse from 3 to 5 dpf, with genes upregulated at 3 dpf becoming downregulated at 5 dpf and vice versa. However, we also find key groups of transcripts and proteins with expression patterns that depend on *chd7* gene dosage regardless of developmental time point. Furthermore, many of these genes that are differentially regulated in MUT relative to HT are associated with neurodevelopmental and transcriptional regulation functions.

### *chd7* indirectly regulates developmental RNA-processing and translation pathways

To define how *chd7* contributes to normal developmental changes in gene expression, we compared transcriptomes from 5 dpf to 3 dpf within each genotype, WT, HT, and MUT. First, we defined DEGs by comparing 5 dpf samples to their corresponding 3 dpf samples and made volcano plots to visualize the distribution of upregulated and downregulated genes across this timeframe (Figure 2A-C). We captured 32,530 RNAs in the WT, HT, and MUT samples, and in all three genotypes more than twice as many genes were up-regulated at 5 dpf than were down-regulated (Figure 2A-C). We then used Ingenuity Pathway Analysis (IPA) to determine canonical pathways that were differentially regulated in each *chd7* genotype (Supplemental Table 7-9). Multiple cell cycle regulatory pathways were similarly altered in all genotypes (Figure 2D-F), indicating that these are independent of *chd7*. We also observed substantial activation of multiple RNA and translational regulation pathways in WT such as nonsense-mediated decay, eukaryotic translation initiation, and EIF2 signaling (Figure 2D) that were not seen in *chd7* HT or MUT, suggesting that *chd7* plays an indirect role in translation by regulating transcript levels of key molecules in these pathways. This is further supported by our observation that in HT and MUT – but not WT – processing of capped intron-containing pre-mRNA was strongly inhibited (Figure 2E, F). This analysis also uncovered significant dysregulation of estrogen signaling from 3 to 5 dpf in *chd7* HT and MUT but not WT (Figure 2D-F), and while hypogonadism in both sexes is commonly seen in CHARGE (Pinto et al., 2005), because our samples were enriched for brain tissue this finding may indicate a role for *chd7* in estrogen-mediated brain development.

**Figure 2.**
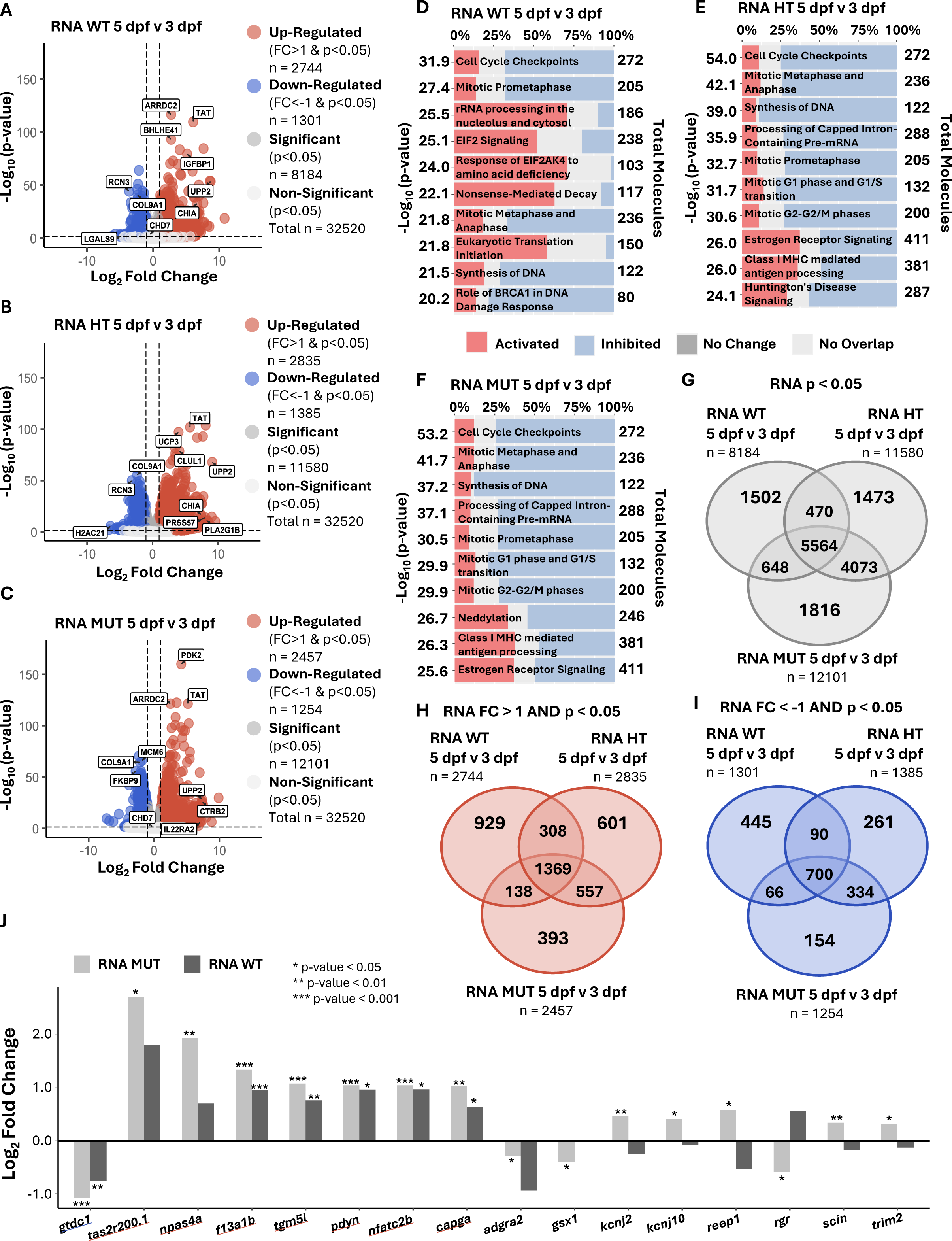
Neural-related transcripts with protein counterparts change developmentally from 3 dpf to 5 dpf due to loss of *chd7.* Volcano plot of Differentially Expressed Genes (DEGs) **A**. 5 dpf Wild Type (WT) compared to 3 dpf WT samples, **B**. 5 dpf Heterozygous (HT) compared to 3 dpf HT samples, and **C**. 5 dpf Homozygous Mutant (MUT) compared to 3 dpf MUT samples. Ingenuity Pathway Analysis (IPA) pathway enrichment by patterns of DEGs from **D**. 5 dpf WT compared to 3 dpf WT samples, **E**. 5 dpf HT compared to 3 dpf HT samples, and **F**. 5 dpf MUT compared to 3 dpf MUT samples. Overlap of DEGs in WT 5 dpf v 3 dpf, HT 5 dpf v 3 dpf, and MUT 5 dpf v 3 dpf **G**. DEGs with p-value < 0.05, **H**. DEGs up-regulated with Fold Change (FC) > 1 and p-value < 0.05, **I**. DEGs down-regulated with FC < 1 and p-value < 0.05. **J**. Log_2_ FC for DEGs selected for most differing FC from **S1.** and all DEGs from overlap with Neuro-GO list **S2.** underlined in red and **S3.** underlined in blue.

We next created Venn diagrams to visualize the differences in developmental changes in gene expression between genotypes. Overall (Figure 2G), we saw more overlap between HT and MUT than between either HT or MUT and WT. We observed a similar pattern when we plotted only genes that were upregulated (Figure 2H) or downregulated (Figure 2I) at 5 dpf. Hypothesizing that important *chd7*-dependent neurodevelopmental genes would have uniquely disrupted patterns of expression from 3 to 5 dpf in *chd7* null mutants, we isolated the MUT DEGs that did not overlap with WT or HT and filtered for neuro-related genes using our Neuro-GO set of genes. We then compared expression levels of these genes in MUT and WT samples (Figure 2J and Figure S1). Many of these neuro-related DEGs were up- or down-regulated in the same direction in WT and MUT, but several were changed in the opposite direction. *Potassium inwardly rectifying channel subfamily J member 2* (*kcnj2*)*, potassium inwardly rectifying channel subfamily J member 10 (kcnj10), receptor accessory protein 1 (reep1), scinderin (scin*), and *tripartite motif containing 2* (*trim2*) are up-regulated in MUT samples and down-regulated in WT samples, while *retinal G protein coupled receptor* (*rgr*) is down-regulated in MUT and up-regulated in WT samples (Figure 2J and Figure S2-3).

Together, these analyses of developmental changes in gene expression reveal that genomic regulation of many developmental processes proceed normally without *chd7,* but neural-related genes, including *kcnj2*, *kcnj10*, *reep1, scin, trim2*, and *rgr* have differing expression patterns in MUT and WT samples and may be required for Chd7-dependent development. Not only that, but loss of *chd7* in the HT and MUT samples results in general inhibition of canonical pathways when compared to WT canonical pathway enrichment.

### Loss of *chd7* in early neurodevelopment causes bidirectional disruption of multiple key neural signaling pathways

To more directly examine how *chd7* regulates gene and protein expression during early neurodevelopment at 3 dpf, we compared RNA and protein levels in HT and MUT samples to WT. We identified a set of 1091 DEGs and 45 DEPs in *chd7* HT and 1257 DEGs and 41 DEPs in *chd7* MUT (Figure 3A-D). Confirming the validity of our CHARGE model, we found that *chd7* RNA and protein were both significantly downregulated in MUT samples. Similar numbers of transcripts and proteins were up- and down-regulated in HT and MUT. We then used Ingenuity Pathway Analysis (IPA) to find canonical pathways with significantly altered enrichment patterns in each genotype (Figure 3E-H and Supplemental Table 10-13). Several pathways were impacted in both HT and MUT at the RNA level, including calcium signaling, striated muscle contraction, and actin cytoskeleton signaling. We did not find pathways significantly altered in both HT and MUT at the protein level, likely because we found many fewer DEPs than DEGs. The only canonical pathway that was disrupted at both the RNA and protein level was SNARE signaling in HT samples. In addition to calcium, SNARE, and actin signaling, we found multiple neural-related molecular pathways were disrupted in HT or MUT at 3 dpf: visual phototransduction, serotonin receptor signaling, synaptogenesis signaling, activation of NMDA receptors, myelination signaling, synaptic long-term potentiation, GABAergic receptor signaling, (Figure 3E-H). Together these impacts indicate broad disruption of neural development in the absence of *chd7*.

**Figure 3.**
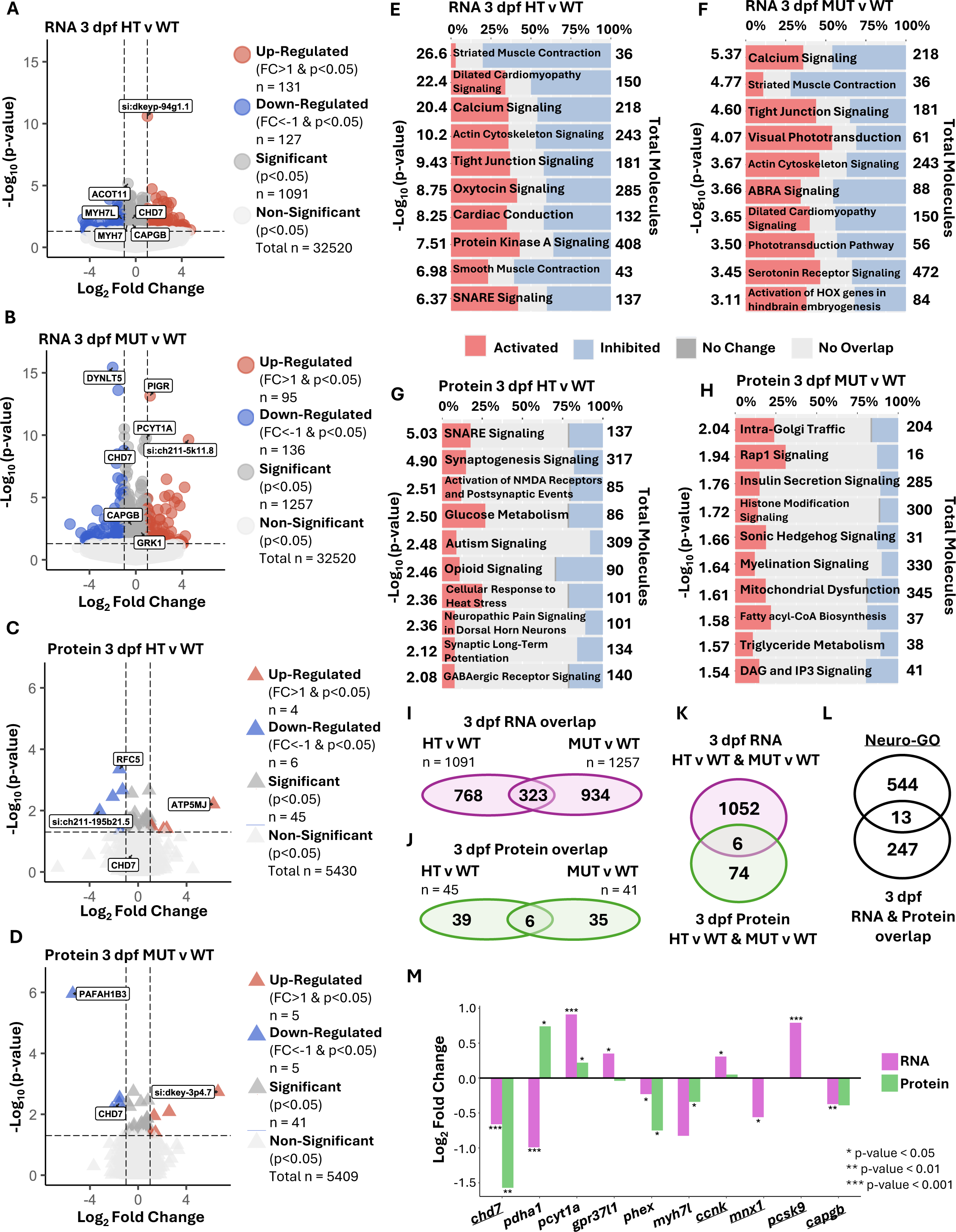
Early neurodevelopmental transcripts with protein counterparts and neural signaling pathways emerge due to loss of *chd7* at 3 dpf. Volcano plot of Differentially Expressed Genes (DEGs) comparing **A**. 3 dpf Heterozygous (HT) compared to 3 dpf Wild Type (WT) samples and **B**. 3 dpf Homozygous Mutant (MUT) compared to 3 dpf WT samples. Volcano plot of Differentially Expressed Proteins (DEPs) comparing **C**. 3 dpf HT compared to 3 dpf WT samples and **D**. 3 dpf MUT compared to 3 dpf WT samples. Ingenuity Pathway Analysis (IPA) pathway enrichment by patterns of DEGs from **E**. 3 dpf HT compared to WT samples and **F**. 3 dpf MUT compared to WT samples. IPA pathway enrichment by patterns of DEPs from **G**. 3 dpf HT compared to WT samples and **H**. 3 dpf MUT compared to WT samples. **I**. Overlap of DEGs with p-value < 0.05 from 3 dpf HT v WT and 3 dpf MUT v WT. **J**. Overlap of DEPs with p-value < 0.05 from 3 dpf HT v WT and 3 dpf MUT v WT. **K**. Overlap of DEGs with p-value < 0.05 from 3 dpf HT v WT and MUT v WT (purple) and DEPs with p-value < 0.05 from 3 dpf HT v WT and MUT v WT (green). **L**. Overlap of 3 dpf DEGs from **I**. and DEPs from **J**. and Neuro-GO list. **M**. Log_2_ Fold Change for all DEGs and DEPs from MUT v WT comparison from overlap in **K**. and selected from overlap in **L**. are underlined.

To identify individual genes that may mediate the effects of Chd7 we created Venn diagrams to highlight genes that were consistently dysregulated at both RNA and protein levels (Figure 3I-L). We found 6 transcripts with protein counterparts that were significantly different (p-value < 0.05) at 3 dpf in either HT or MUT (Figure 3K). None of these 6 overlaps with the Neuro-GO list (Figure S4). We also found 13 neuro-related transcripts with protein counterparts that were significantly differentially expressed (p-value < 0.05) at either the transcript or protein level (Figure 3L and Figure S5). We then plotted both transcript and protein expression for these genes and found that most were consistently up- or down-regulated as both RNA and protein, with the exception of *pyruvate dehydrogenase E1 subunit alpha 1a* (*pdha1a*), for which the transcript is down-regulated, and protein is up-regulated (Figure 3M). *chd7, phosphate regulating endopeptidase X-linked (phex), myosin heavy chain 7-like (myh7l),* and *capping actin protein gelsolin like b* (*capgb*) were all down-regulated at both the RNA and protein level, while *phosphate cytidylyltransferase 1A choline* (*pcyt1a*) was up-regulated at both the RNA and protein level (Figure 3M).

These data reinforce previous findings that RNA and protein levels do not always correlate (Nie et al., 2006), and that *chd7* has bidirectional effects on gene expression (Huang et al., 2025; Jamadagni et al., 2021; Stathopoulou et al., 2023; Yao et al., 2020). Furthermore, we find that *chd7* regulates multiple neural signaling pathways and that *capgb*, due to its consistent dysregulation at RNA and protein levels and across *chd7* genotypes, may be a key effector of *chd7* in neural development.

### Loss of *chd7* disrupts RNA processing and ROBO receptor signaling in 5 dpf zebrafish

To identify Chd7 target genes that may mediate zebrafish CHARGE model behavioral phenotypes that emerge at 5 dpf, we compared 5 dpf transcriptomes and proteomes data from *chd7* HT and MUT to WT. We identified a set of 3263 DEGs and 56 DEPs in *chd7* HT and 2491 DEGs and 460 DEPs in *chd7* MUT (Figure 4A-D). Homozygous loss of *chd7* at 5 dpf caused two times more DEGs to be down-regulated than up-regulated (680 down vs 290 up) (Figure 4B), and we saw a similar pattern at the protein level, with 88 downregulated and 65 upregulated proteins. We next used IPA to uncover molecular pathways that are dysregulated in the absence of *chd7* based on DEGs (Figure 4E-F and Supplemental Table 14-15) and DEPs (Figure 4G-H and Supplemental Table 16-17). At the RNA level, most altered canonical pathways were inhibited rather than activated in *chd7* HT and MUT, and multiple RNA regulatory pathways were disrupted in both HT and MUT, including EIF2 signaling, nonsense-mediated decay, and eukaryotic translation initiation, elongation, and termination (Figure 4E-F). We also observed decreases in ROBO receptor signaling in both HT and MUT, indicating potential for axon guidance defects. At the protein level, only one canonical pathway was disrupted in both HT and MUT: processing of capped intron-containing pre-mRNA (Figure 4G-H), providing further indication of disruptions to RNA processing with loss of *chd7*.

**Figure 4.**
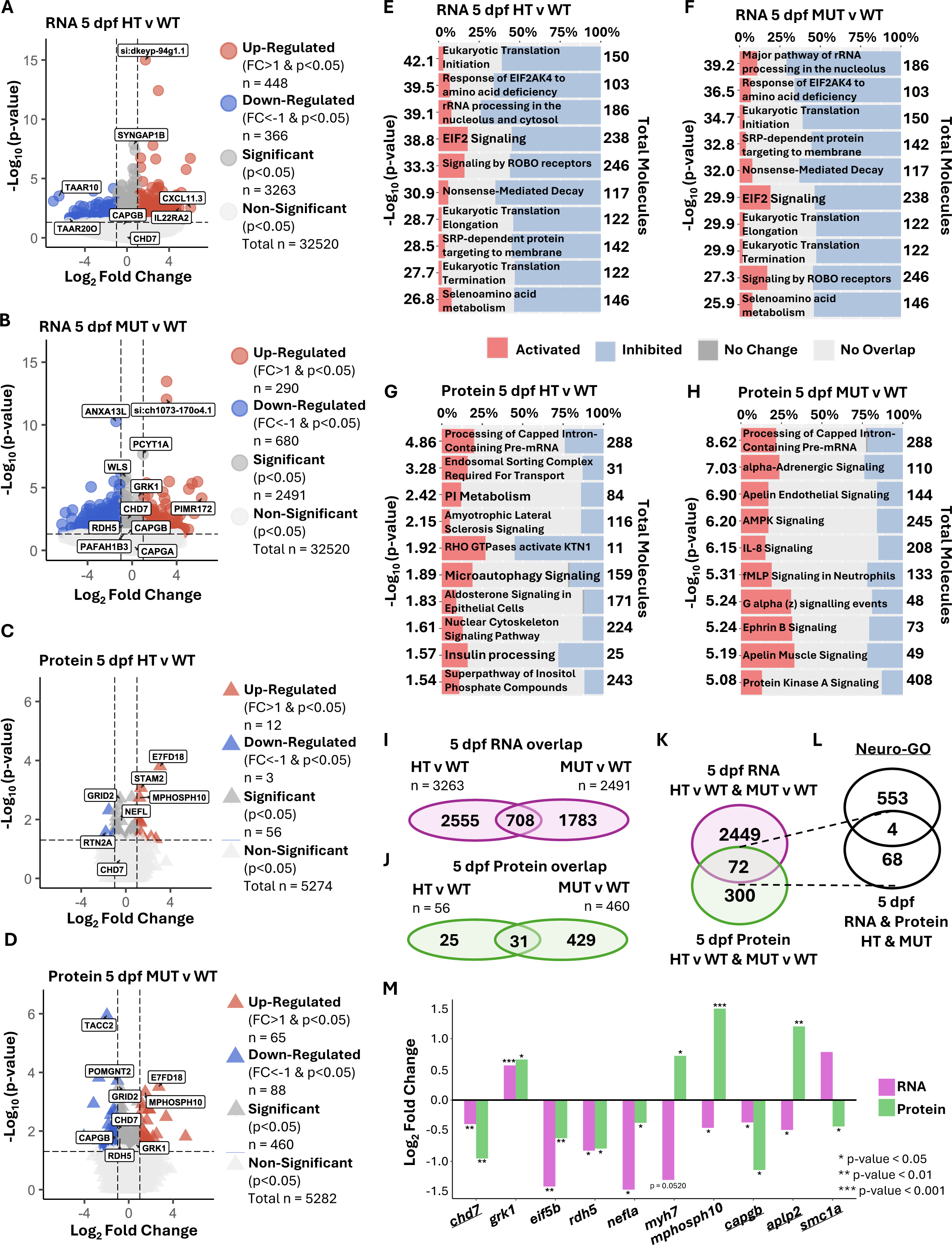
At 5 dpf down-regulated transcripts with protein counterparts and inhibition of pathways emerge. Volcano plot of Differentially Expressed Genes (DEGs) comparing **A**. 5 dpf Heterozygous (HT) compared to 5 dpf Wild Type (WT) samples and **B**. 5 dpf Homozygous Mutant (MUT) compared to 5 dpf WT samples. Volcano plot of Differentially Abundant Proteins (DEPs) comparing **C**. 5 dpf HT compared to 5 dpf WT samples and **D**. 5 dpf MUT compared to 5 dpf WT samples. Ingenuity Pathway Analysis (IPA) pathway enrichment by patterns of DEGs from **E**. 5 dpf HT compared to WT samples and **F**. 5 dpf MUT compared to WT samples. Ingenuity Pathway Analysis (IPA) pathway enrichment by patterns of DEPs from **G**. 5 dpf HT compared to WT samples and **H**. 5 dpf MUT compared to WT samples. **I**. Overlap of DEGs with p-value < 0.05 from 5 dpf HT v WT and DEGs with p-value < 0.05 from 5 dpf MUT v WT. **J**. Overlap of DEPs with p-value < 0.05 from 5 dpf HT v WT and 5 dpf MUT v WT. **K**. Overlap of DEGs with p-value < 0.05 from 5 dpf HT v WT and MUT v WT (purple) and DEPs with p-value < 0.05 from 5 dpf HT v WT and MUT v WT (green). **L**. Overlap of 5 dpf DEGs and DEPs from **K**. and Neuro-GO list. **M**. Log_2_ FC for selected DEGs and DEPs from MUT v WT comparison from overlap in **K**. and overlap in **L**. underlined.

We next created Venn diagrams to visualize the shared transcripts with protein counterparts that were dysregulated in *chd7* HT and MUT samples (Figure 4I-L). We found 72 transcripts with protein counterparts that were significantly different (p-value < 0.05) at 5 dpf in either HT or MUT (Figure 4K and Figure S6). Of those 72 genes, 3 were up-regulated and 39 were down-regulated in both transcript and protein. Furthermore, 4 of the 72 are canonically neuro-related (Figure 4L): *chd7, capping actin protein gelsolin like b (capgb), amyloid beta (A4) precursor-like protein 2 (aplp2)*, and *structural maintenance of chromosomes 1A (smc1a)*. Of these, *chd7* and *capgb* were consistently downregulated at the RNA and protein levels, while *aplp2* and *smc1a* had opposing transcript and protein expression patterns (underlined in Figure 4M). 6 more of the 72 genes were selected and plotted by prioritizing significance and presence in the MUT sample: *eukaryotic translation initiation factor 5B (eif5b), retinol dehydrogenase 5 (rdh5), neurofilament light chain a (nefla), G protein coupled receptor kinase 1 (grk1), myh7,* and *mphosph10*. Of these, *eif5b, rdh5,* and *nefla* were consistently downregulated at the RNA and protein levels while *grk1* was consistently upregulated. These unbiased analyses further confirm the validity of our zebrafish CHARGE model in showing that *chd7* is strongly downregulated, and they strengthen the possibility that *capgb*, which was consistently downregulated at 3 and 5 dpf, may be an important molecular mediator of Chd7.

### Identification of *chd7*-dependent upstream regulators, functions, and potential mediators of CHARGE phenotypes

To reveal *chd7*-dependent molecular processes from consistent gene expression patterns in HT and MUT samples, we used IPA to further integrate our transcriptomic and proteomic analyses from 3 and 5 dpf. First, we used patterns of DEGs and DEPs to define enriched canonical pathways in HT and MUT (Figure 5A-B and Supplemental Table 18-19). These canonical pathways, upstream regulators, and diseases and functions analyses consider patterns of up- and down-regulation, and by also integrating multiple timepoints and sample comparisons we were able to identify trends between conditions. DEG data showed consistent disruption of multiple cardiac and neural pathways across genotypes and timepoints: apelin cardiomyocyte, cardiac conduction, cardiac hypertrophy signaling, sensory processing of sound by inner and outer hair cells of the cochlea, calcium signaling, acetylcholine receptor signaling, ROBO-SLIT signaling, and SNARE signaling. DEP data revealed many fewer altered pathways, most likely due to the smaller set of DEPs identified compared to DEGs (Figure 5A). In 5 dpf MUT samples, however, we also found inhibition of several neural pathways such as CREB signaling in neurons, synaptogenesis signaling, glutaminergic receptor signaling, serotonin receptor signaling, and oxytocin in brain (Figure 5B). Next, we identified upstream regulators using IPA by patterns of DEGs and DEPs in HT and MUT samples from 3 and 5 dpf (Figure 5C-D and Supplemental Table 20-21). At 3 dpf most upstream regulators were inhibited while being activated at 5 dpf based on transcript data (Figure 5C). Protein data showed that GABA signaling was disrupted at 5 dpf, along with other key neural factors including RB1 and beta-estradiol (Figure 5D). Finally, our analysis of diseases and functions with IPA (Figure 5E-F and Supplemental Table 22-23) revealed general inhibition at 3 dpf and activation at 5 dpf in both *chd7* HT and MUT (Figure 5E). Several CHARGE-related functions were altered, such as sensory system development, growth failure, disease of retina, and seizure disorder. Together these findings paint a picture in which loss of *chd7* causes widespread molecular dysfunction that differs at the transcript and protein levels and shifts during development.

**Figure 5.**
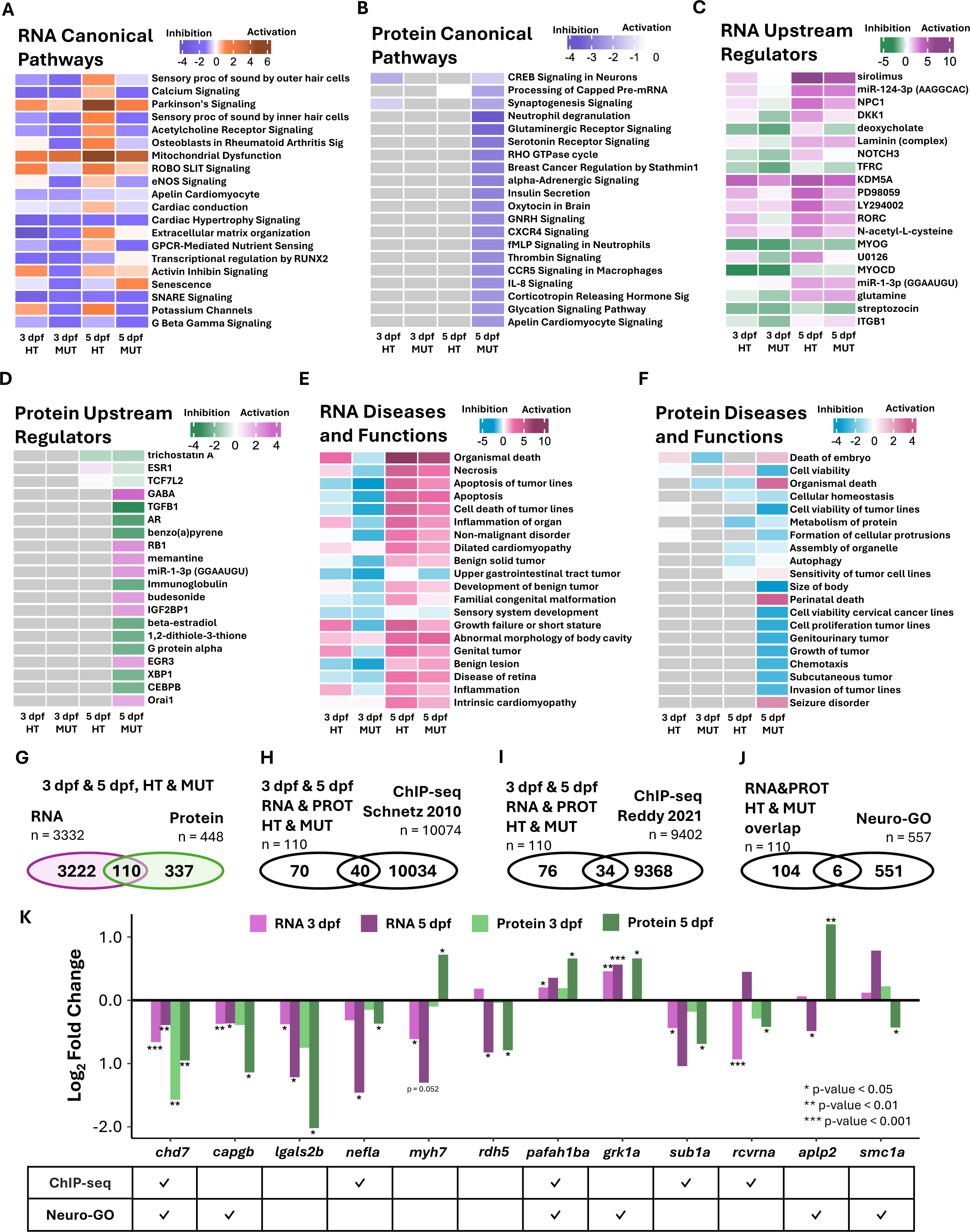
Transcript and protein expression patterns inform downstream pathway, regulator, and functional analysis. Ingenuity Pathway Analysis (IPA) canonical pathway enrichment by patterns of **A**. DEGs with p-value < 0.05 from 3 dpf and 5 dpf HT v WT and MUT v WT comparisons and **B**. DEPs with p-value < 0.05 from 3 dpf and 5 dpf HT v WT and MUT v WT comparisons. IPA upstream regulator enrichment by patterns of **C**. DEGs with p-value < 0.05 from 3 dpf and 5 dpf HT v WT and MUT v WT comparisons and **D**. DEPs with p-value < 0.05 from 3 dpf and 5 dpf HT v WT and MUT v WT comparisons. IPA diseases and functions enrichment by patterns of **E**. DEGs with p-value < 0.05 from 3 dpf and 5 dpf HT v WT and MUT v WT comparisons and **F**. DEPs with p-value < 0.05 from 3 dpf and 5 dpf HT v WT and MUT v WT comparisons. **G**. Overlap of DEGs with p-value < 0.05 3 dpf and 5 dpf HT v WT and MUT v WT comparisons and DEPs with p-value < 0.05 from HT v WT and MUT v WT comparisons. **H**. Overlap of DEGs and DEPs from **G**. with public ChIP-seq data from Schnetz 2010. **I**. Overlap of DEGs and DEPs from **G**. with public ChIP-seq data from Reddy 2021. **J**. Overlap of DEGs and DEPs from **G**. with Neuro-GO list. **K**. Log2 FC of selected candidate DEGs and DEPs from MUT v WT comparison plotted at 3 dpf and 5 dpf, and table below noting presence in ChIP-seq or Neuro-GO list.

The complexity of molecular changes that result from *chd7* loss of function (Figures 1-5) makes it challenging to define individual factors that contribute directly to CHARGE phenotypes, as they most likely emerge from this collection of molecular changes. However, our data sets highlight some consistent patterns across genotypes and timepoints, which support an effort to isolate key genes that may mediate the effects of *chd7* on neural development and behavior. We hypothesized that the most likely candidate genes would be dysregulated at 3 and 5 dpf, in HT and MUT, and at the RNA and protein levels, and would be targets of *chd7*-mediated chromatin accessibility. First, we found 110 genes with protein counterparts that were significantly dysregulated (p < 0.05) in the 3 or 5 dpf sample (Figure 5G and Figure S7). Next, we cross-checked those 110 genes with publicly available ChIP-seq data from mouse embryonic stem cells (Schnetz et al., 2010) and cerebellar granule neurons (Reddy et al., 2021) to narrow in on genes *chd7* may regulate by binding within 3000 bp of a transcription start site (TSS) (Figure 5H-I and Figure S8-9). Finally, we compared these genes with our list neurodevelopmental genes generated from GO Accession terms and Ensembl (Neuro-GO) (Figure 5J). These analyses uncovered 10 candidate genes (Figure 5K) plus *chd7*, which was significantly (p-value < 0.05) down-regulated in all samples, binds within 3000 bp of its own TSS, and is canonically neuro-related. *Capping actin protein gelsolin like b* (*capgb*) was also down-regulated in all samples and is canonically neuro-related. *Lectin galactoside-binding soluble 2* (*lgals2*) was down-regulated in all samples and has been found to be expressed in the zebrafish visual system at 60 hpf (Thijssen et al., 2006). *Neurofilament light chain a* (*nefla*) was also down-regulated in all samples, CHD7 binds within 3000 bp of its TSS, and its paralog *neflb* is known to regulate neuron apoptosis (Wang et al., 2019). In contrast, myosin heavy chain 7 (*myh7*) has opposing transcript and protein expression and is not canonically neuro-related. However, *myh7* is known to be involved in actin filament binding and cardiac development (Hesaraki et al., 2022) and thus may contribute to *chd7*-dependent cardiac development. Similarly, *retinol dehydrogenase 5* (*rdh5*) transcript and protein are significantly downregulated at 5 dpf, and although it is not canonically neuro-related it is known to be involved in retinal development, and variants of *rdh5* in humans cause delayed dark adaption (Yamamoto et al., 1999). Finally, *platelet-activating factor acetylhydrolase 1b* (*pafah1b*) is up-regulated in all samples, CHD7 binds within 3000 bp of its TSS, and it is canonically neuro-related.

In summary, we reveal 6 candidate molecular mediators of *chd7* whose dysregulation may cause CHARGE model phenotypes. Of the 6, there is Chip-seq data indicating that *nefla* and *pafah1b* are directly regulated by Chd7, and *capgb* and *pafah1b* are canonically neuro-related while *lgals2, nefla, myh7,* and *rdh5* have been shown previously to be involved in or expressed during nervous system development.

### Knockdown of candidate genes causes CHARGE-related behavioral and morphological phenotypes

To determine if dysregulation of our candidate genes causes behavioral and/or morphological phenotypes consistent with our zebrafish CHARGE model (Hodorovich et al., 2023), we used the CRISPR-Cas9 system to knock down each gene using two gRNAs targeting different sites in the gene (Table 1). gRNAs were complexed with Cas9 protein and injected into one-cell stage zebrafish embryos. We raised injected embryos to 5 dpf and recorded, tracked, and analyzed behavioral responses of these F0 crispants to acoustic and visual stimuli. We also documented any observed morphological defects. All injected embryos were screened for gene edits, and those in which one or both target sites were edited were included in the analyses (Figure S10), with injected unedited embryos serving as controls (Figure S12). Neither gRNA for *pafah1b* or *lgals2* induced edits (Figure S10-11), so these became our injected controls, and we focused on the remaining 4 candidate genes, *capgb, nefla, rdh5, and myh7*.

**Table 1.**
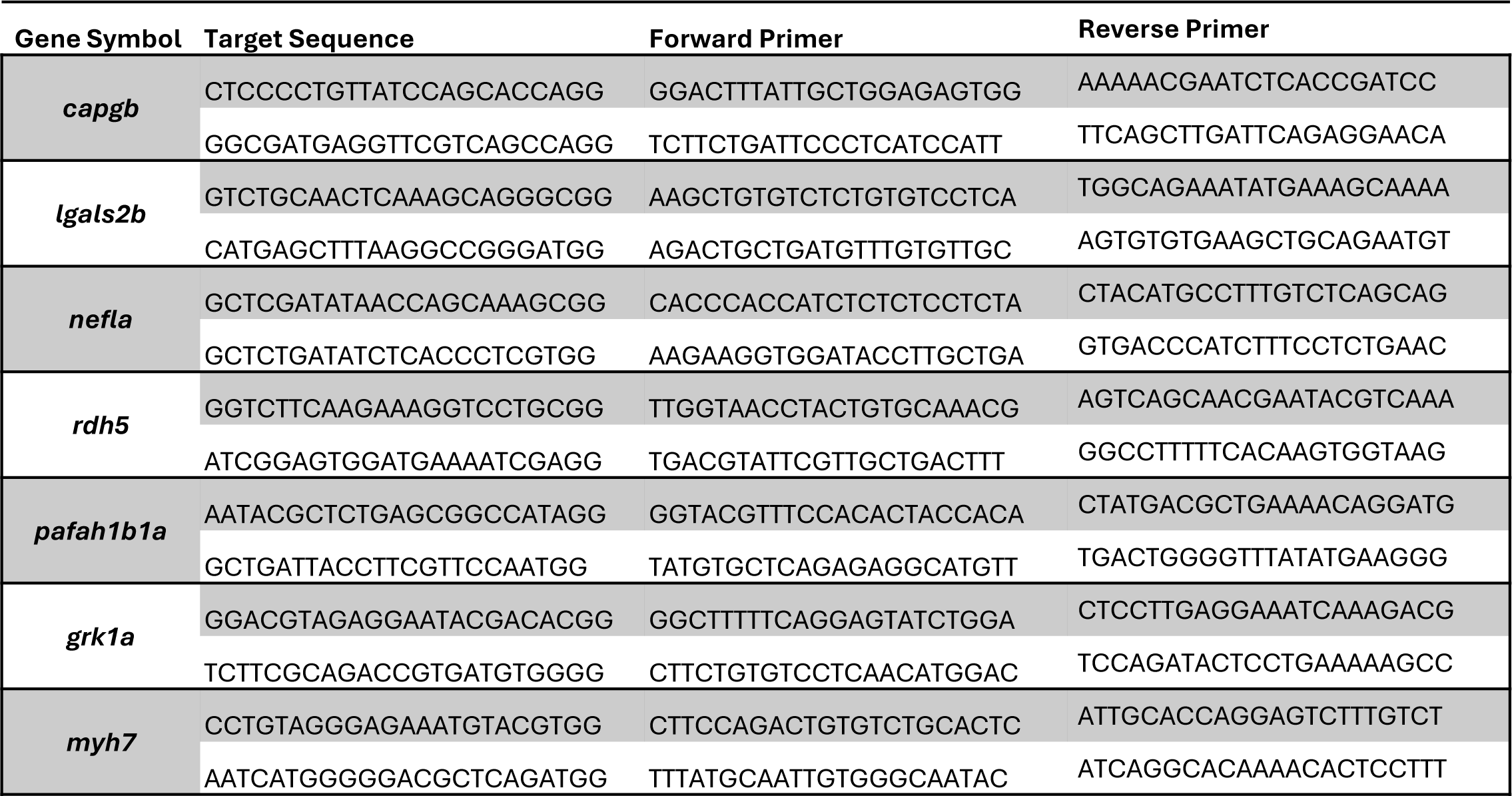
Candidate gene symbol with corresponding gRNA target sequence and primers made using CHOPCHOP.

We tested if sensory processing was affected by knockdown of candidate genes by recording responses of 5 dpf larvae to acoustic stimuli of increasing intensity (Figure 6A-D). We found that knockdown of candidate genes *capgb, nefla, rdh5, and myh7* did not affect Short-Latency C-bend (SLC) response frequency (Figure 6A-B). However, Long-Latency C-bend (LLC) response frequency was reduced upon knockdown of *capgb, nefla,* or *rdh5* compared to injected controls (Figure 6C-D). This phenocopies *chd7* mutant larvae, which also display normal SLC and reduced LLC responses (Hodorovich et al., 2023).

**Figure 6.**
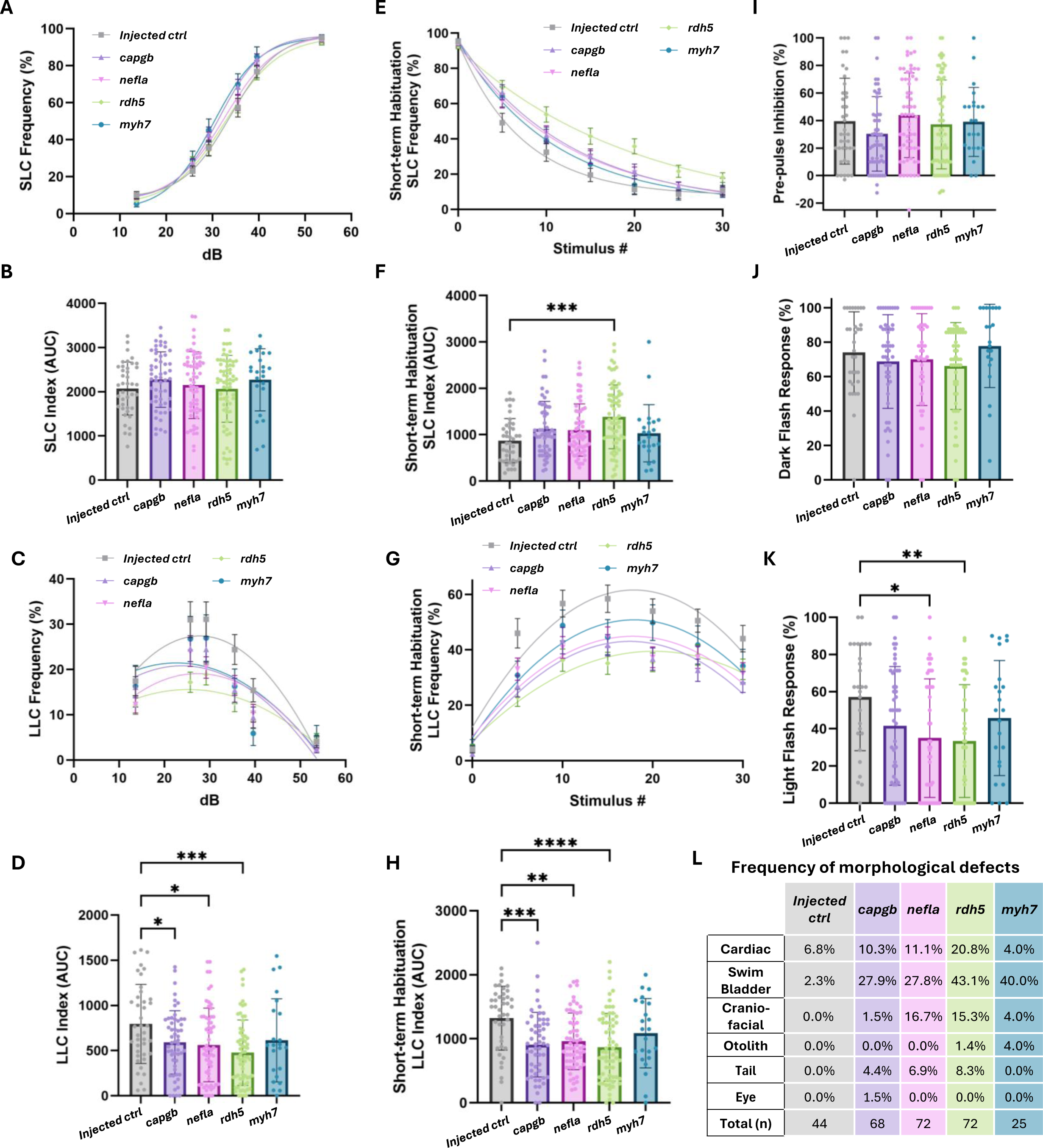
Knockdown of candidate genes causes phenocopy of CHARGE model phenotypes **A**. Short-Latency C-bend (SLC) frequency plotted against acoustic stimulus intensity for each group of larvae fit with a nonlinear regression sigmoidal dose response curve. **B**. Area under the curve (AUC) plotted for each individual larva’s SLC response frequency. **C**. Long-Latency C-bend (LLC) frequency plotted against acoustic stimulus intensity for each group of larvae fit with a nonlinear regression second order polynomial (quadratic) curve. **D**. AUC plotted for each individual larva’s LLC response frequency. **E**. Short-term Habituation (STH) SLC frequency plotted against time for each group of larvae fit with nonlinear regression one-phase decay curve. **F.** STH AUC for each individual larva’s SLC response frequency. **G.** STH LLC frequency plotted against time for each group of larvae fit with a nonlinear regression second order polynomial (quadratic) curve. **H.** STH AUC for each individual larva’s LLC response frequency. **I.** Pre-pulse Inhibition percent. **J.** Dark flash response frequency. **K**. Light flash response frequency. **L.** Frequency of morphological defects observed for each group of larvae. (Bar graphs: MeanL±LStandard Deviation, Line graphs: MeanL±LSEM, Ordinary one-way ANOVA with Dunnetts’s multiple comparisons, * p-value < 0.05, ** p-value < 0.01, *** p-value < 0.001, **** p-value < 0.0001).

We next tested candidate gene crispants for proper habituation to repeated strong acoustic stimulation, a simple form of non-associative learning. Habituation of SLCs was reduced by knockdown of *rdh5* (Figure 6E-F). We also found that LLCs, which are normally increased during habituation, were reduced in *capgb, nefla,* and *rdh5* crispants compared to injected controls (Figure 6G-H). These phenotypes are opposite to those of *chd7* mutants, which show faster SLC habituation and increased LLCs following repeated acoustic stimulation (Hodorovich et al., 2023). We also tested another form of startle modulation, pre-pulse inhibition (PPI), and found that knockdown of any of our candidate genes had no effect on PPI (Figure 6I).

Next, we measured responses to visual stimuli following knockdown of each candidate gene (Figure 6J-K). All crispants displayed normal O-bend response frequency when given a series of 10 dark flash stimuli (Figure 6J). However, knockdown of *nefla* or *rdh5* caused a decrease in light flash response frequency compared to injected controls (Figure 6K). Finally, we assessed the presence of morphological defects that are seen in *chd7* mutants, including cardiac, swim bladder, craniofacial, and otolith defects (Hodorovich et al., 2023). Knockdown of each of the 4 candidate genes caused an increase in the frequency of observed defects compared to injected controls (Figure 6L), with knockdown of *rdh5* resulting in the highest frequency of morphological phenotypes with 20.8% having cardiac defects, 43.1% swim bladder defects, 15.3% craniofacial defects, and 1.4% otolith defects (Figure 6L). Otolith defects are very rare in clutches of wildtype embryos but are the most penetrant phenotype in *chd7* mutants (Hodorovich et al., 2023). We saw otolith defects in both *rdh5* and *myh7* crispants, indicating a potential role for both genes in mediating *chd7*-dependent otolith formation.

In summary, knockdown of *capgb, nefla,* or *rdh5* causes behavioral defects in response to acoustic and visual stimuli similar to what is seen in *chd7* mutants. Additionally, loss of any of our 4 candidate genes results in morphological defects that are also observed in *chd7* mutants. Together, these findings provide evidence that *capgb*, *nefla*, *rdh5*, and *myh7* may be novel molecular mediators of CHD7 that contribute to CHARGE-related developmental and behavioral phenotypes.

## Discussion

The chromatin remodeler CHD7 is critical for the development of many tissues, as evidenced by the widespread deficits seen in CHARGE patients, which often include complex behavioral challenges including Autism Spectrum Disorder, sensory processing defects, altered pain sensation, aggression, obsessive compulsive disorder, and anxiety (Blake et al., 2005; Edwards et al., 1995; Hartshorne et al., 2005, 2017). The molecular pathophysiology of these neurobehavioral symptoms is not well understood, and so in this study we sought to uncover and functionally validate molecular mediators of Chd7-dependent neurodevelopment using an established zebrafish CHARGE model that recapitulates many CHARGE features (Hodorovich et al., 2023). By integrating bulk transcriptomic and proteomic data from two developmental timepoints in *chd7* wildtypes, heterozygotes, and homozygous mutants, we reveal gene expression patterns and canonical pathways that drive *chd7*-dependent neurodevelopment. Finally, using CRISPR/Cas9 we confirmed that multiple candidate genes play roles in zebrafish CHARGE-related behavioral and morphological phenotypes.

CHD7 is a master transcriptional regulator that can enhance or repress transcription (Schulz, Wehner, et al., 2014). Our data reinforces this concept in that loss of *chd7* does not result in global up- or down-regulation of transcripts and proteins, but rather we observed that *chd7* has bidirectional effects on developmental gene expression. These findings are consistent with transcriptomic analyses of 5 dpf mutant *chd7* zebrafish larval brains (Jamadagni et al., 2021), *Chd7* null mouse embryonic stem cells (ESCs) (Yao et al., 2020), conditional knock-out of *Chd7* in cardiac mouse embryo tissue (Stathopoulou et al., 2023), and *Chd7* mutant mouse embryo forebrain tissues (Huang et al., 2025). In contrast, CHD8, a closely related member of the CHD family of chromatin remodelers, seems to primarily enhance transcription as homozygous knockout of *CHD8* in neurons derived from human induced pluripotent stem cells (hIPSCs) results in markedly more downregulated genes than upregulated genes (Haddad Derafshi et al., 2022). The dual role for *chd7* in both promoting and repressing gene expression adds to the complexity of determining the molecular targets whose dysregulation leads to CHARGE symptoms upon *chd7* loss of function.

Proper recruitment of transcription factors and chromatin remodelers like CHD7 is crucial for regulating gene expression. CHD7 is recruited to enhancers at specific H3K4me1 and H3K4me3 histone modifications and promotes H3K27ac histone modifications by histone acetylases (Lettieri et al., 2021; Reddy et al., 2021). We found enrichment of histone modification signaling (Figure 3H) and activation of upstream regulator *kdm5a* (Figure 5C), a histone demethylase that regulates H3K4me3 marks (El Hayek et al., 2020). Similarly, mutations in both *CHD7* and *KDM5A* histone modifying genes are associated with congenital heart disease (Zaidi et al., 2013), and variants in *KDM5A* have been associated with Autism Spectrum Disorder (El Hayek et al., 2020). This is consistent with CHARGE patients having behavioral symptoms including Autistic-like behaviors and heart defects (Hartshorne et al., 2005; Verloes, 2005). Not only that, but CHD7 also interacts with CHD8 (Batsukh et al., 2010), and variants in CHD8 are strongly associated with Autism Spectrum Disorder (Weissberg and Elliott, 2021). Additionally, mutations in *KDM6A,* are associated with Kabuki syndrome (Van Laarhoven et al., 2015), which has clinical diagnostic criteria that overlap with CHARGE syndrome (Schulz, Freese, et al., 2014). These findings suggest that loss of *CHD7* may give rise to some CHARGE behavioral and morphological phenotypes through its role in regulating histone modifications via histone demethylase KDM5A.

The primary focus of this study was on the molecular basis of neurodevelopmental deficits caused by loss of *chd7*. Based on patterns of DEGs and DEPs in *chd7* HT and MUT samples, we found enrichment of many neurodevelopmental pathways. One prominently inhibited pathway was calcium signaling (Figure 3E-F, 5A). Calcium is a key second messenger with many cellular roles including neuronal signaling and plasticity (Rosenberg and Spitzer, 2011), muscle contraction (Fearnley et al., 2011), and apoptosis (Clapham, 2007). We also found inhibition of GABAergic receptor signaling (Figure 3A) and activation of upstream regulator GABA (Figure 5D). It has previously been found that *chd7* is required for GABAergic neuron development via *paqr3b* regulation in 5 dpf zebrafish (Jamadagni et al., 2021). While we did not observe significant dysregulation of *paqr3b* in our datasets, our findings further support the role of GABAergic inhibition in *chd7*-mediated neural development and function. We also found impacts of loss of *chd7* on excitatory signaling, with dysregulation of NMDA receptors and postsynaptic events (Figure 3G), synaptic long-term potentiation (Figure 3G), and down-regulation of glutamate receptor ionotropic delta 2 (Grid2) protein (Figure 4C-D). Consistent with these findings, *kismet*, the *Drosophila* orthologue of *CHD7* and *CHD8*, has been shown to be important for localization of postsynaptic glutamate receptors and synaptic transmission (Ghosh et al., 2014). GRID2 does not bind glutamate (Araki et al., 1993) but alternatively binds d-serine and glycine (Naur et al., 2007) and has been found to be important for NMDA receptor function (Kumagai et al., 2014) and synaptic plasticity in mouse models (Kakegawa et al., 2011). GRID2 is also expressed in cerebellar Purkinje cells (Araki et al., 1993), where it has roles in synapse organization (Matsuda et al., 2010). This aligns with our observation of disrupted synaptogenesis signaling in *chd7* HT at 3 dpf (Figure 3G). Previously, conditional knockout of *Chd7* in mouse models was found to induce cerebellar hypoplasia and distinct cerebellar foliation anomalies leading to motor delay and coordination deficits (Whittaker, Kasah, et al., 2017). Thus, one possibility is that loss of *CHD7* alters cerebellar expression of GRID2, leading to reduced synapse formation and altered excitability, contributing to cerebellar hypoplasia seen in some CHARGE patients (Shiohama et al., 2019).

Our analyses uncovered several additional neurodevelopmental processes affected by loss of *chd7*, including axon guidance, myelination, cytoskeletal regulation, and SNARE signaling. Signaling by ROBO receptors (Figure 4E-F) and ROBO-SLIT signaling (Figure 5A), which are crucial for axon guidance throughout central nervous system development (Brose et al., 1999), were significantly altered in *chd7* HT and MUT. This is not the first link between *chd7* and ROBO function, as previous transcriptional analysis identified down-regulation of genes associated with ROBO-SLIT signaling in Chd7 mutant mouse heart tissue (Payne et al., 2015). We also observed inhibition of myelination signaling (Figure 3H), which is consistent with earlier work showing that inactivation of Chd7 causes defects in oligodendrocyte myelination and that Chd7 is required for oligodendrocyte remyelination *in vitro* (He et al., 2016). Our data also highlights multiple cytoskeletal regulatory pathways that were disrupted by loss of *chd7*, including actin cytoskeleton signaling, which we discuss below, and downregulation of neurofilament light chain protein a (*nefla*) (Figure 4M, 5K). *nefla* is orthologous to human *NEFL* and provides structural support to axons (Yuan et al., 2017). Finally, SNARE signaling was strongly inhibited at both the RNA and protein levels (Figure 3E, 3G, 5A), suggesting that disruptions to the fundamental processes of neurotransmitter release and receptor trafficking may contribute to CHARGE symptoms. Most of these neurodevelopmental processes were disrupted more strongly at 3 dpf than 5 dpf, suggesting that *chd7* largely functions to promote the proper formation of neural circuits rather than their function. Together, the data from our zebrafish model provide confirmation of many previous findings regarding *chd7*’s conserved roles in neural development while also setting a foundation for further investigation of the novel mechanisms our analyses have revealed.

Several actin-related processes were affected by loss of *chd7* including actin cytoskeleton signaling (Figure 3E-F), striated muscle contraction (Figure 3E-F), ABRA (Actin-Binding Rho Activating Protein) signaling (Figure 3F), RHO GTPase activity (Figure 4G, 5B), and down-regulation of capping protein actin filament gelsolin-like b (*capgb*) (Figure 3M,4M). *Capgb* is orthologous to human *CAPG*, an actin-regulatory protein that is part of the gelsolin family that serves a crucial role in the organization of the actin cytoskeleton (Johnston et al., 1990) and is known to be part of the TYRO protein tyrosine kinase-binding protein (TYROBP) causal network in microglia (Zhang et al., 2013). Proper regulation of actin dynamics is crucial for many cellular processes in which *chd7* is also implicated, particularly in the nervous system, including neurogenesis (Pacheco and Gallo, 2016), cell migration (Okuno et al., 2017), inner ear development (Tilney et al., 1980), and glial development (Sepp and Auld, 2003). These key roles in CHARGE-related mechanisms and the persistent disruption of *capgb* expression we observed suggest that additional investigation of the relationship between *chd7* and *capgb* is warranted.

Deficits in ear morphology, auditory processing, and deafness are the most common symptoms seen in CHARGE patients (Morgan et al., 1993) and are observed in CHARGE models (Bajpai et al., 2010; Bosman et al., 2005; Patten et al., 2012). For example, in mouse models haploinsufficiency of *Chd7* and *Sox2* results in reduced otic cell proliferation, severe malformations of semicircular canals, and shortened cochleae with ectopic hair cells (Gao et al., 2024). Conditional deletion of Chd7 in mouse also results in cochlear hypoplasia and complete absence of the semicircular canals and cristae (Hurd et al., 2010). Our zebrafish model also recapitulates some of these defects, including altered morphology of otoliths, calcium crystal structures in the otic vesicles that are vibrated by auditory stimuli and activate hair cells (Baeza-Loya and Raible, 2023), which is the most penetrant morphological phenotype we observed (Hodorovich et al., 2023). Perhaps one of the most striking findings of the current study is that CRISPR/Cas9 knockdown of candidate mediators *rdh5* and *myh7* causes similar defects in otolith morphology (Figure 6L). The penetrance of these defects is much lower than what is seen in *chd7* HT and MUT larvae, however, indicating that otolith morphology defects are likely driven by combined dysregulation of multiple genes with *chd7* loss of function.

We previously found that *chd7* mutant larvae have specific deficits in responding to auditory stimuli with Long-Latency C-bends (LLCs) but not Short-Latency C-bends (SLCs) (Hodorovich et al., 2023). This LLC deficit is independent of otolith defects, however, as both *chd7* mutants with and without otolith defects display reduced LLC frequency (Hodorovich et al., 2023). This indicates that additional auditory processing defects drive this behavioral deficit. Our current multi-omic analyses further support this conclusion, as canonical pathway analysis of *chd7* HT and MUT showed inhibition of sensory processing of sound by inner and outer hair cells (Figure 5A) and inhibition of sensory system development (Figure 5E). And based on consistent dysregulated expression across timepoints in *chd7* HT and MUT, we identified and directly tested four potential mediators of *chd7*-dependent neurodevelopment, three of which when knocked down with CRISPR/Cas9 phenocopied the LLC deficit seen in *chd7* mutants (Figure 6C-D). These data provide strong evidence that *capgb*, *nefla*, and *rdh5* contribute to *chd7*’s regulation of auditory function. That knockdown of just one of these, *rdh5*, resulted in a weakly penetrant otolith defect is consistent with the conclusion that downstream auditory processing is altered by loss of *chd7*.

*capgb* and *nefla*, through their functions in regulating the cytoskeleton can be linked with multiple aspects of neural development, but *rdh5* is perhaps a more surprising regulator of auditory responses. Rdh5 functions to catalyze the final step in the production of 11-cis retinaldehyde, the universal chromophore in visual pigments (Jang et al., 2001) and contributes to the metabolism of retinoic acid (RA) (Mao et al., 2023), an important morphogen for hindbrain development, where the neural circuits that drive LLC and SLC responses are located. RA signaling has also previously been linked to inner ear development in CHARGE mouse models (Micucci et al., 2014). Furthermore, Hox genes are downstream of RA (reviewed in Nolte et al., 2019), and our current pathway analyses of *chd7* mutants uncovered disruption of Hox genes in hindbrain embryogenesis (Figure 3F). While *capgb*, *nefla*, and *rdh5* crispants phenocopy the *chd7* LLC deficit following non-habituating auditory stimuli, these crispants also showed reduced LLCs during habituation (Figure 6H), opposite to the increased LLCs seen during habituation in *chd7* mutants (Hodorovich et al., 2023). This difference is most likely because knockout of each of our candidate genes disrupts just one rather than the many pathways in combination that occurs with loss of *chd7*. Also, crispants are mosaic such that not every cell is expected to harbor a loss-of-function mutation in the target gene, so we may not expect to see complete phenocopy of *chd7* mutants in the F0 generation even if that gene is a direct mediator of *chd7*’s effects on the LLC circuit.

Eye defects and visual impairments are additional common characteristics of CHARGE patients that have been investigated in CHARGE animal models (Krueger and Morris, 2022). Chd7 expression in the retina and retinal defects have been observed and characterized in zebrafish and mouse CHARGE models (Krueger et al., 2023). Our current molecular pathway analyses of zebrafish *chd7* mutants also reflect disrupted visual processing, as visual phototransduction (Figure 3F) and disease of retina (Figure 5E) were significantly enriched. We also found that knockdown of candidate molecular mediators *nefla* and *rdh5* causes a decrease in behavioral responses to light flashes with no significant difference in dark flash response frequency (Figure 6J-K). *chd7* mutants, however, display normal light flash responses and reduced dark flash responses (Hodorovich et al., 2023). This difference could again reflect the mosaicism of the crispants and that just a single *chd7* mediator was targeted. While it is also possible that *nefla* and *rdh5* act independently of *chd7* in the visual system, these findings support further investigation of how these pathways contribute to defects in vision caused by loss of *chd7*.

In summary, we have found that loss of *chd7* causes broad dysregulation of neurodevelopmental, sensory system, and gene regulatory processes. From this multi-omic analysis several molecular mediators of CHARGE model behavioral phenotypes have emerged. In future work it will be important to measure the combinatorial effects the candidate effector genes *capgb, nefla,* and *rdh5* have on auditory and visual responses to determine if they function in overlapping gene networks. Another key consideration is that *chd7* has tissue-specific roles in regulating gene transcription (Williams et al., 2024), so tissue-specific manipulation of *capgb, nefla,* and *rdh5* would help to illuminate their roles in different parts of the nervous system. Finally, our data highlights many additional genes and pathways that were beyond the scope of this investigation that are likely contributors to CHARGE pathology that are worthy of investigation, such as Grid2, estrogen signaling, and Slit-ROBO-mediated axon guidance. Overall, by integrating transcriptomic and proteomic analysis across two key developmental timepoints, we have found that potential molecular targets of Chd7 *capgb*, *nefla*, and *rdh5* mediate CHARGE model behavioral phenotypes. These findings will help to define new genes and pathways regulated by Chd7 to may be potential therapeutic targets to alleviate the neurobehavioral aspects of CHARGE syndrome.

## Materials and Methods

### *Danio rerio* husbandry and maintenance

All animal use and procedures were approved by and in accordance with the North Carolina State University Institutional Animal Care and Use Committee (IACUC) guidelines. All *chd7* Wild Type (WT), Heterozygous (HT), and Homozygous mutant (MUT), and all candidate gene crispants used were of the *Tüpfel long fin* (TLF) background. The TLF strain originated from Zebrafish International Resource Center (ZIRC) stocks. Adult zebrafish were housed in 5Lfish/L density under a 14 h:10 h light:dark cycle at ∼28°C, and were fed rotifers, *artemia* brine shrimp (Brine Shrimp Direct), and GEMMA micro 300 (Skretting).

To generate embryos for larval testing, male and female pairs were placed in mating boxes (Aquaneering) containing system water and artificial grass. 1–2 h into the subsequent light cycle of the following day, embryos were collected and placed into petri dishes containing 1× E3 embryo media. Embryos were sorted for fertilization under a dissecting scope at ∼6Lh post fertilization (hpf) and placed into 10 cm petri dishes with *n*L≤L65. All embryos were reared in a temperature-controlled incubator at 29°C on a 14 h:10 h light:dark cycle. Each day until testing, a 50%–75% media change was performed.

### Zebrafish model of CHARGE syndrome

In this study we use a previously characterized zebrafish model of CHARGE syndrome (Hodorovich et al., 2023). CRISPR-Cas9 gene editing induced a 7 bp frameshift deletion that leads to a premature stop codon in exon 9 of chd7.

### Dissections

Larvae were anesthetized at 3 days post fertilization (dpf) or 5 dpf zebrafish with ice cold E3. Dissections were performed under a dissecting microscope with scalpel and forceps. A scalpel was used to cut diagonally to include all brain tissue and exclude yolk. Head tissue was rinsed in 1 X PBS and then placed in RNA later (RNA) or 1 X PBS (protein) for long term storage. Head tissue in RNA later was placed on ice for 20 mins, 4C for 24 hours, and then -20C for long term storage based on manufacturer protocol. The corresponding tail tissue was placed in Methanol for subsequent genotyping. Genotyping was done by prepping DNA from tail tissue using Hotshot DNA extraction. Then that DNA was used in a PCR reaction with GoTaq DNA polymerase and primer pair forward: GATGATGAGCCCTTCAACCCAG and reverse: CAGATGGTTTGAGAACGATTGA. PCR reactions were visualized using gel electrophoresis with 50 bp ladder where wild type bands are 132 bp and mutant bands are 125 bp.

### Transcriptomics

#### RNA isolation

Once genotypes were known, head tissue was pooled based on genotype, 40 heads for 3 dpf and 20 heads for 5 dpf, using a glass pasteur pipette into a 1.5 mL tube then stored at -20C until ready for isolation. For RNA isolation New England Biolabs Monarch Total RNA miniprep kit and protocol were used. Follow “Part 1: Sample disruption and homogenization” and “Part 2: RNA binding and elution” for tissue up to 10mg. RNA concentration was determined using an Implen NanoPhotometer and then sent to the North Carolina State University Genomic Sciences Laboratory (GSL) for 250 bp paired end sequencing on the Illumina Novaseq.

#### RNA-sequencing analysis

Demultiplexed reads were analyzed using the NCSU Bioinformatics Resource Center (BRC) computing cluster and SLURM commands can be found at https://github.com/melody-create. Quality Control was done with FastQC. Adapter trimming and quality filtering was done with Fastp. An indexed reference was built using GRCz11.fna and GRCz11.gtf input ensembl.org/info/data/ftp/index.html. Paired end sample reads were aligned with STAR aligner. Differential expression analysis of un-normalized counts was done with DESeq2. All RNA-sequencing p-values calculated by Wald test in DESeq2.

### Proteomics

#### Protein isolation

Once genotypes were known, head tissue was pooled based on genotype, 20 heads for 3 dpf and 10 heads for 5 dpf, using a glass pasteur pipette into a 1.5 mL tube then stored at -20C until ready for isolation. For protein isolation tissue was suspended in a 100 µL solution of 50 mM ammonium bicarbonate (pH 8.0) containing 1% Sodium deoxycholate (SDC). Samples were lysed by probe sonication via 2 pulses at 20 seconds per pulse at a 20% amplitude setting. Cellular debris was removed via centrifugation at 10,000 RPM for 5 minutes at 4C. Supernatant was retained with pipette on ice and assessed for protein quantification by bicinchoninic acid (BCA) assay using the Pierce BCA Protein Assay Kit. The standard curve was used to determine protein concentration of each unknown sample.

### Materials

The following materials were purchased from Thermo Fisher Scientific (Wilmington, DE): 1 M tris hydrochloride solution pH 7.5, 1 M tris hydrochloride solution pH 8, sodium chloride, ammonium bicarbonate (ABC), Pierce™ Mass Spec Grade trypsin, Pierce™ BCA Protein Assay, Pierce™ Quantitative Colorimetric Peptide Assay, LC/MS grade water, LC/MS grade acetonitrile, LC/MS grade formic acid, and Vivicon 30 kD molecular weight cutoff filters. Urea, dithiothreitol (DTT), and iodoacetamide (IAA) were purchased from Bio-Rad (Hercules, CA). Sodium deoxycholate (SDC) and calcium chloride, were purchased from MilliporeSigma (St. Louis, MO).

### Filter Aided Sample Preparation (FASP)

Isolated protein was reconstituted into 50 mM ABC, 1% SDC and a 200 uL aliquot precipitated and rinsed with 800 uL ice-cold acetone. Protein was allowed to precipitate for 30 min at -20 °C. After centrifugation acetone was decanted and removed entirely by vacuum evaporation. Rinsed protein was reconstituted again in 50 mM ABC, 1% SDC and the concentration measured by Pierce™ BCA Protein Assay. A 200 uL volume of each sample containing about 27 µg protein was incubated with 15 μL of 50 mM DTT at 56 °C for 30 min. Samples were transferred to Vivicon 30 kD filters and washed with 8 M urea in 0.1M TRIS-HCl pH 8. Cysteines were alkylated with 64 μL of 200 mM iodoacetamide and incubation at room temperature in the dark for 1 hr. Samples were rinsed 3 times with 2 M urea, 10 mM CaCl_2_ in 0.1M Tris-HCl pH 8, then 3 times with 0.1M Tris-HCl pH 7. All rinsates were discarded. Trypsin protease solution was added to reach a 1:25::trypsin:protein ratio, and samples were incubated overnight at 37 °C. Amounts of recovered peptides were quantified by Pierce™ Quantitative Colorimetric Peptide Assay. Samples were evaporated to dryness in a vacuum concentrator and reconstituted in 98% water, 2% acetonitrile, 0.1% formic acid to reach a protein concentration of 0.5 µg/µL.

### LC-MS/MS analysis

A 2 µL injection was analyzed by reversed phase nano-liquid chromatography-mass spectrometry (nano-LC-MS/MS) using an Easy-Nano-1200 nanoLC system (Thermo Scientific, San Jose, CA, USA) interfaced with an Orbitrap Exploris 480 (Thermo Scientific) Mass Spectrometer. The ‘trap and elute’ configuration consisted of a 0.075 mm × 20 mm C_18_ trap column with particle size of 3 µm (Thermo Scientific Accclaim PepMap^TM^ 100, Part # 164946) in line with a 0.075 mm × 250 mm C_18_ nanoLC analytical column with particle size of 2 µm (Thermo Scientific PepMap^TM^, Part # ES902). Peptides were eluted using a solvent gradient of water containing 2% acetonitrile, 0.1% formic acid (MPA) and acetonitrile containing 20% water, 0.1% formic acid (MPB). MPB was held at 5% for 2 min, increased to 25% over 100 min, increased to 40% over 15 min, increased to 95% in 1 min, and was held at 95% for 13 min. Mass spectrometer parameters were set as follows: 2.0 kV positive ion mode spray voltage, ion transfer tube temperature of 275 °C, master scan cycle time of 3 s, *m/z* scan range of 375 to 1,600 at 120,000 _FWHM_ resolving power (at *m/z* 200), 300% normalized AGC Target, 120 ms maximum MS^1^ injection time, RF lens of 40%, 15,000 _FWHM_ resolving power (at *m/z* 200) for data-dependent MS^2^ scans, 0.7 *m/z* isolation window, 30% normalized HCD collision energy, 100% normalized AGC Target, automated maximum injection time and dynamic exclusion applied for 60 s periods.

### Data Interrogation

Raw nanoLC-MS/MS files were processed with Proteome Discoverer 2.5 software (PD, Thermo Scientific, San Jose, CA) using *Danio rerio* (Taxon 7955) protein databases obtained from Swiss-Prot (3,274 sequences) and TrEMBL (83,386 sequences). A custom contaminants database was included in the searches to identify presence of human keratin and reagent enzyme peptides. Trypsin was designated as the cleaveage reagent with hydrolysis sites at the c-terminus of lysine and arginine. A label-free workflow was employed to obtain protein abundance values. The SEQUEST HT search node was set up with the following parameters: maximum of 3 missed cleavage sites; minimum peptide length of 6 amino acids; 5 ppm precursor mass tolerance; 0.02 Da fragment mass tolerance; maximum of 4 dynamic modifications per peptide, which were oxidation of methionine, n-terminal acetylation, and methionine loss; static carbamidomethylation of cysteine. Peptides were validated by Percolator with q-value set to 0.05 and strict false discovery rate (FDR) set to 0.01. Protein abundances were calculated using all peptides with normalization to total peptide amount. No scaling was performed. Hypothesis testing on replicate comparisons used ANOVA.

### Transcriptomics and Proteomics Integration Analyses

All transcriptomic and proteomic data was plotted using R packages and tools and the R code can be found at https://github.com/melody-create. Principal component analysis was done using DESeq2 (Love et al., 2014) for transcriptomic data and Proteome Discoverer for proteomic data. Heatmaps were made using Complex Heatmap (Gu et al., 2016) (Gu, 2022). RNAs and proteins from heatmap slices were analyzed for gene ontology using the PANTHER 19.0 Overrepresentation Test (Released 20240807), DOI: 10.5281/zenodo.15066566 Released 2025-03-16, Danio rerio (all genes in database), GO biological process complete, Fisher’s Exact test, False Discovery Rate correction (https://geneontology.org/). Volcano plots were made using Enhanced Volcano (Blighe, K, S Rana, and M Lewis, 2018). Raw data from Qiagen Ingenuity Pathway Analysis (IPA) canonical pathways, upstream regulators, and diseases and functions was plotted using ggplot and Complex Heatmap.

### Quiagen Ingenuity Pathway Analysis (IPA)

Canonical pathway analysis, upstream regulator analysis, and diseases and functions analysis were done using Quiagen Ingenuity Pathway Analysis (IPA). Fold change and p-value data were imported and filtered by p value < 0.05.

### Chip-seq data

Previous Chip-seq datasets were downloaded from NCBI NIH Gene Expression Omnibus (GEO): GSM558674 (Schnetz et al., 2010) and GSE164360 (Reddy et al., 2021) and analyzed in R using ChIP seeker (Wang et al., 2022) and R code can be found at https://github.com/melody-create.

### Crispant generation and analysis

Gene targets were selected based on omics data integration. Guide RNA (gRNA) target sites and primers were selected using the online web tool CHOPCHOP (version 3). Flanking primers were ordered from Integrated DNA Technologies (IDT). Tissue was collected from Wild type Tupelo Long Fin (TLF) strain background zebrafish DNA extracted and target sites sequenced using primer pairs stated above to match target sequence provided from CHOPCHOP to actual strain sequence. Once gRNA targets sites were confirmed CRISPR RNA (crRNA) was ordered from IDT.

CRISPR/Cas9 injection cocktails were prepared by combining crRNA, tracrRNA, Cas9 and phenol red. Wild type Tupelo Long Fin (TLF) strain background zebrafish were bred and their fresh embryos collected. 1 nanoliter of injection mix was injected into one cell stage embryos. The embryos were raised to 5 dpf and phenotyping was performed by capturing morphology and behavior. After phenotyping, each larva was genotyped by extracting DNA from tissue, PCR reaction using primers flanking the target sequence, and resulting reactions were analyzed by gel electrophoresis. There was said to be an edit by CRISPR/Cas9 if the resulting banding pattern contained extra bands, smears, smaller bands, or larger bands when compared to wild type uninjected controls. Injected control larvae were prepared the same as above and had no edits when analyzed by gel electrophoresis or by sanger sequencing.

### Behavior testing and analysis

5 dpf zebrafish larvae were loaded onto a 6x6 grid with wells filled with 1 X E3 media. The grid is attached to a programmable acoustic shaker which delivers sound stimuli of varying intensities directly to the grid. Also part of the behavior rig is a programmable light source which delivers ambient light, light flash (whole field illumination), and dark flash (whole field darkness). There is a camera above which captures images at 1000 frames per second (fps), with 1/1600 shutter speed at 640 X 640 pixel resolution. These parameters are set using Photron Fastcam Viewer (PFV) software. Event timer software was used to trigger the camera to capture the set amount of frames after a stimulus. 1000 frames are captured after visual stimuli and 120 frames are captured after acoustic stimuli. The assays were 10X light flash, 10X dark flash, 60X startle, 10X prepulse inhibition, 30X short term habituation. Once images are captured, FLOTE software was used for unbiased tracking of each larval zebrafish based on head and tail location. Tracked data was analyzed for specific behaviors using FLOTE batch analysis software. This output was then converted to an excel spreadsheet cleaned and compiled for analysis. Data was loaded into Graphpad Prism 10.4.2 where average, standard deviation, and count were calculated and plotted as a dot plot. The area under the curve was calculated for each larval behavior output and plotted as a startle index as a show all points bar plot.

### Morphology

Once behavior was captured each 5 dpf zebrafish larva was placed in an individual well of a 24 well plate and tricaine was added. Larvae were looked at under a dissecting microscope at 3X magnification. Each larva was looked at for morphological abnormalities in the otolith, swim bladder, eye, heart, tail, craniofacial, and body.

## Acknowledgments

We would like to thank the NCSU Genomic Science Library (GSL) for performing quality control and sequencing of the transcriptomic data. We would like to thank the CHHE for providing a discount on our sequencing services. We would like to thank Leonard Collins, Ying Xi, and Taufika Islam Williams at METRIC NCSU for performing Filter Aided Sample Preparation, LC-MS/MS analysis, and Data Interrogation of the proteomics data. We would like to thank Stack Overflow and its users for providing many answers to coding questions. We would like to thank Sureni Sumathipala, Kimberly Charron, and Alexandra Venuto for their support. We would like to thank Emily Costanzo for her help auditing citations.

## Data and resource availability

All data is retained locally and is available upon request. Raw transcriptome and proteome data is available on the Dryad repository. All code used for analysis is available at https://github.com/melody-create.

## Author contributions

*Project conceptualization*: K.C.M and M.B.H. *Methodology*: K.C.M and M.B.H.. *Investigation*: M.B.H., D.R.R., R.A.B, D.C.C, and K.C.M. *Data curation*: M.B.H. *Writing*: M.B.H. and K.C.M. *Writing – review & editing*: M.B.H., D.R.R., R.A.B, D.C.C, and K.C.M. *Supervision*: K.C.M. *Funding acquisition*: K.C.M.

