## Supplementary material for "Multi-omic analyses identify molecular targets of Chd7 that mediate CHARGE syndrome model phenotypes": Figure S1

Supplemental Figure 1. List of gene symbol overlap between Neuro-GO list and RNA p < 0.05 MUT 5 dpf v 3 dpf from Figure 2G.

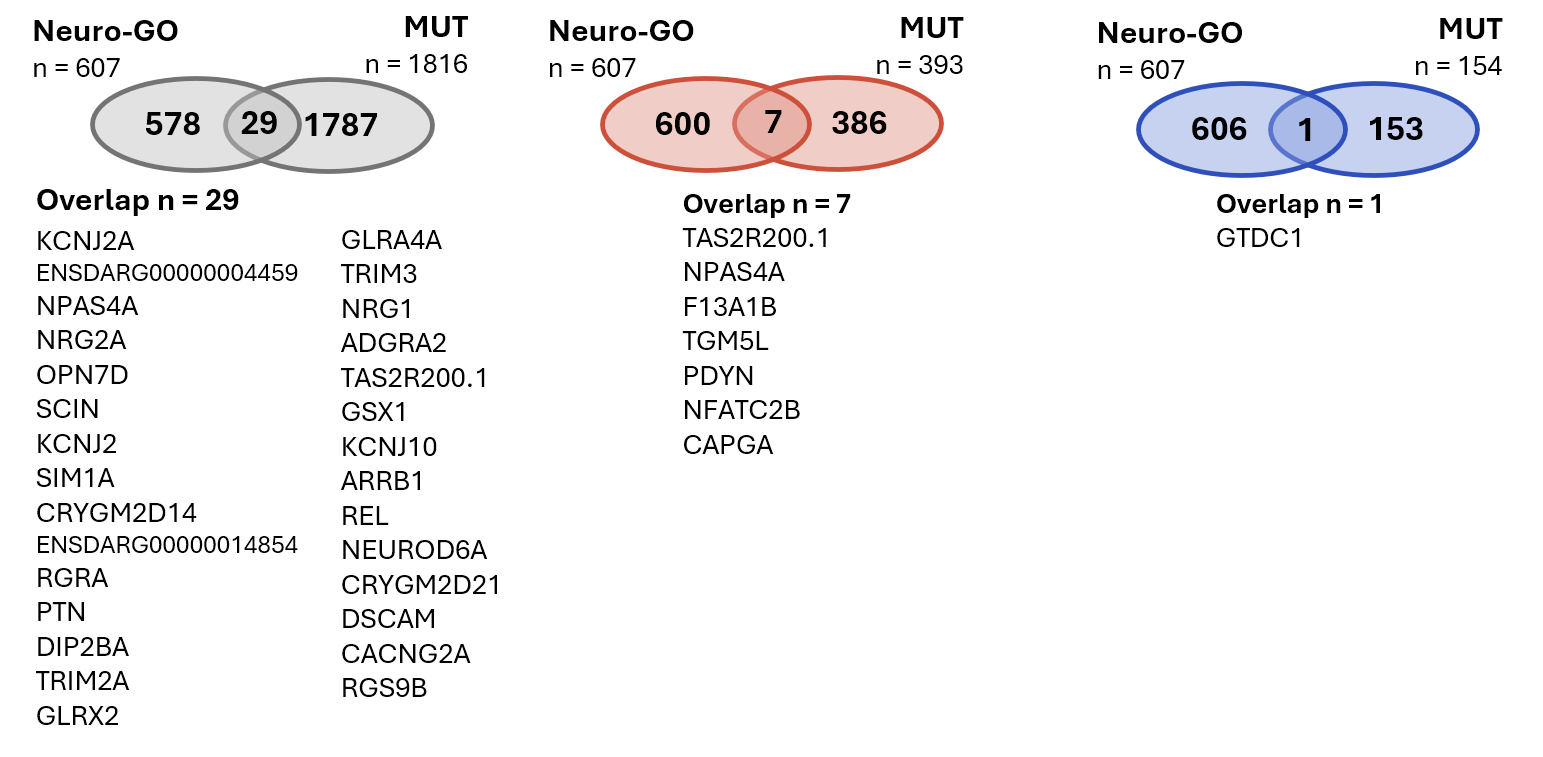

Supplemental Figure 2. List of gene symbol overlap between Neuro-GO list and RNA FC > 1 AND p < 0.05 MUT 5 dpf v 3 dpf from Figure 2H.

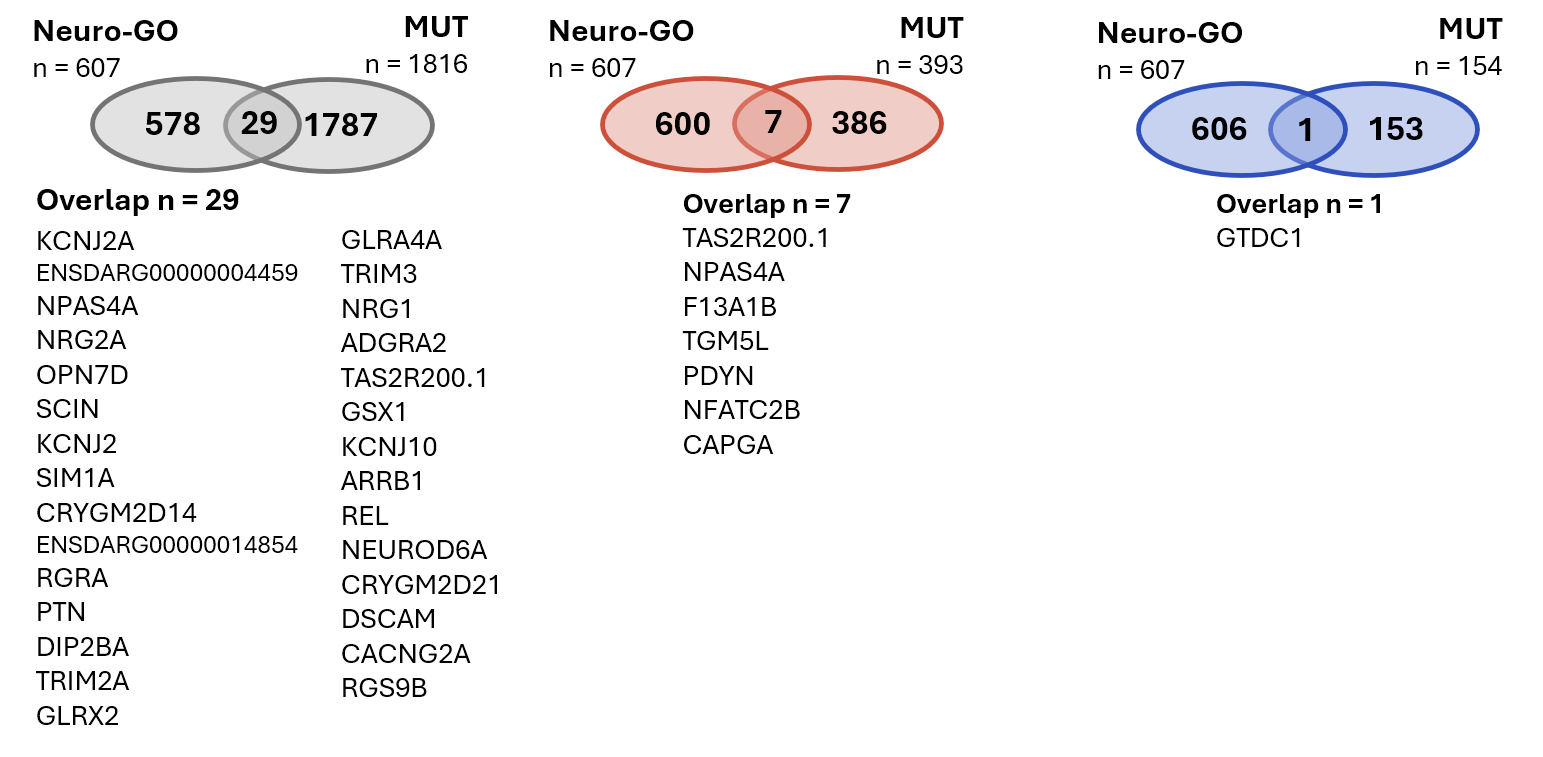

Supplemental Figure 3. List of gene symbol overlap between Neuro-GO list and RNA FC < -1 AND p < 0.05 MUT 5 dpf v 3 dpf from Figure 2I.

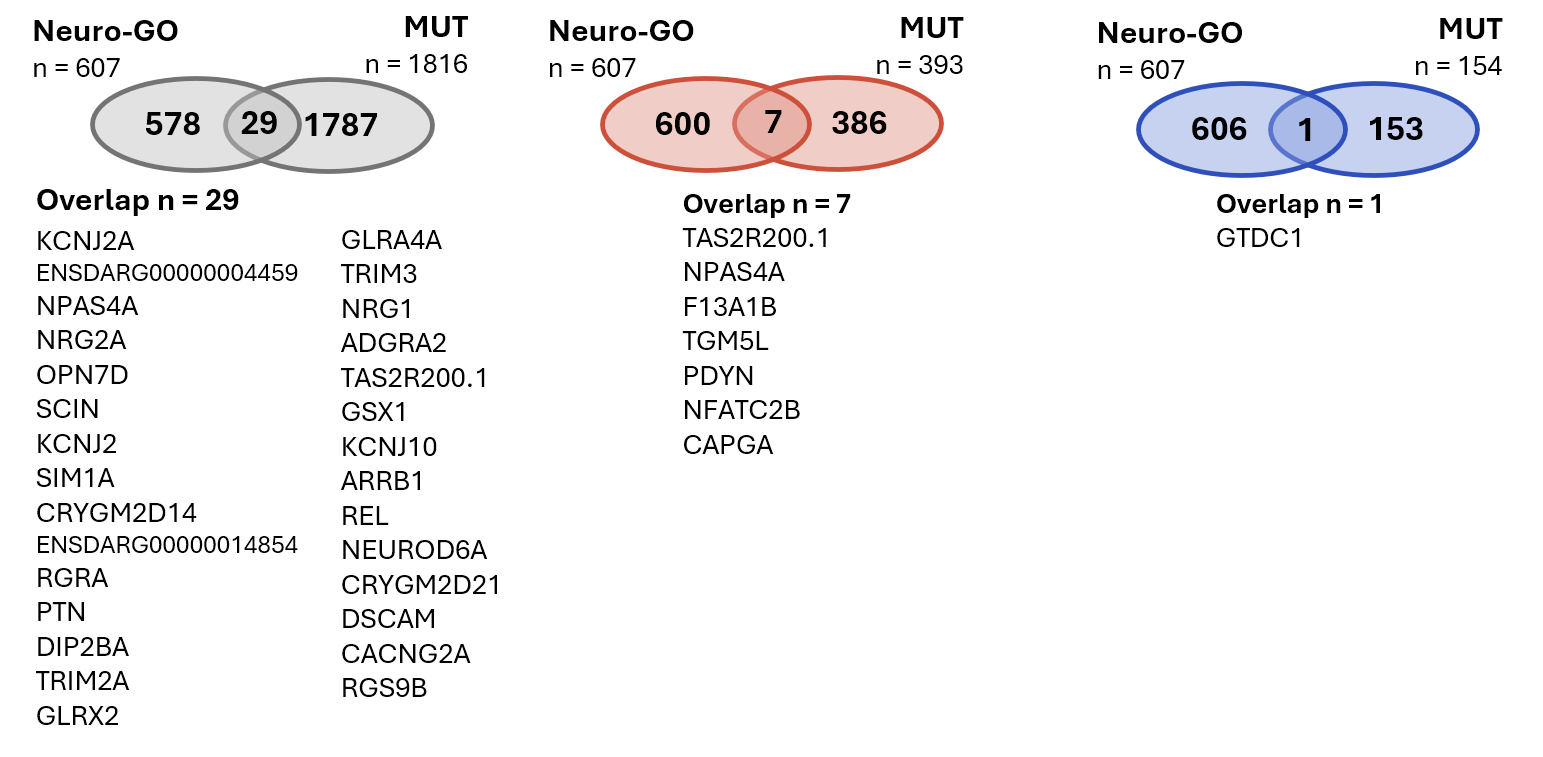

Supplemental Figure 4. List of gene symbol overlap between 3 dpf RNA HT v WT & MUT v WT and 3 dpf Protein HT v WT & MUT v WT from Figure 3K. Showing no overlap between Neuro-GO list and 3 dpf RNA & Protein HT & MUT which is the same comparison made in Figure 4L.

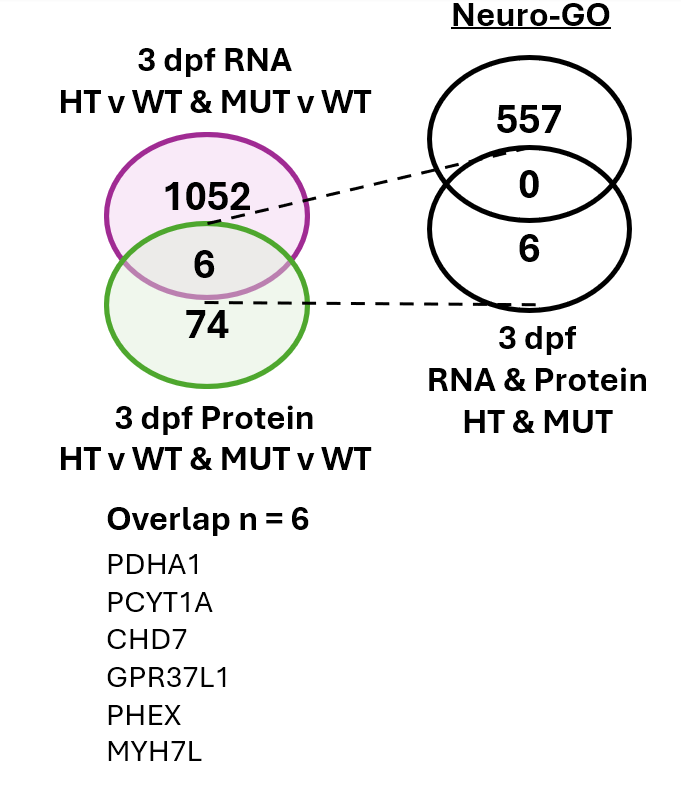

Supplemental Figure 5. List of gene symbol overlap between Neuro-GO list and 3 dpf RNA & Protein overlap from Figure 3L.

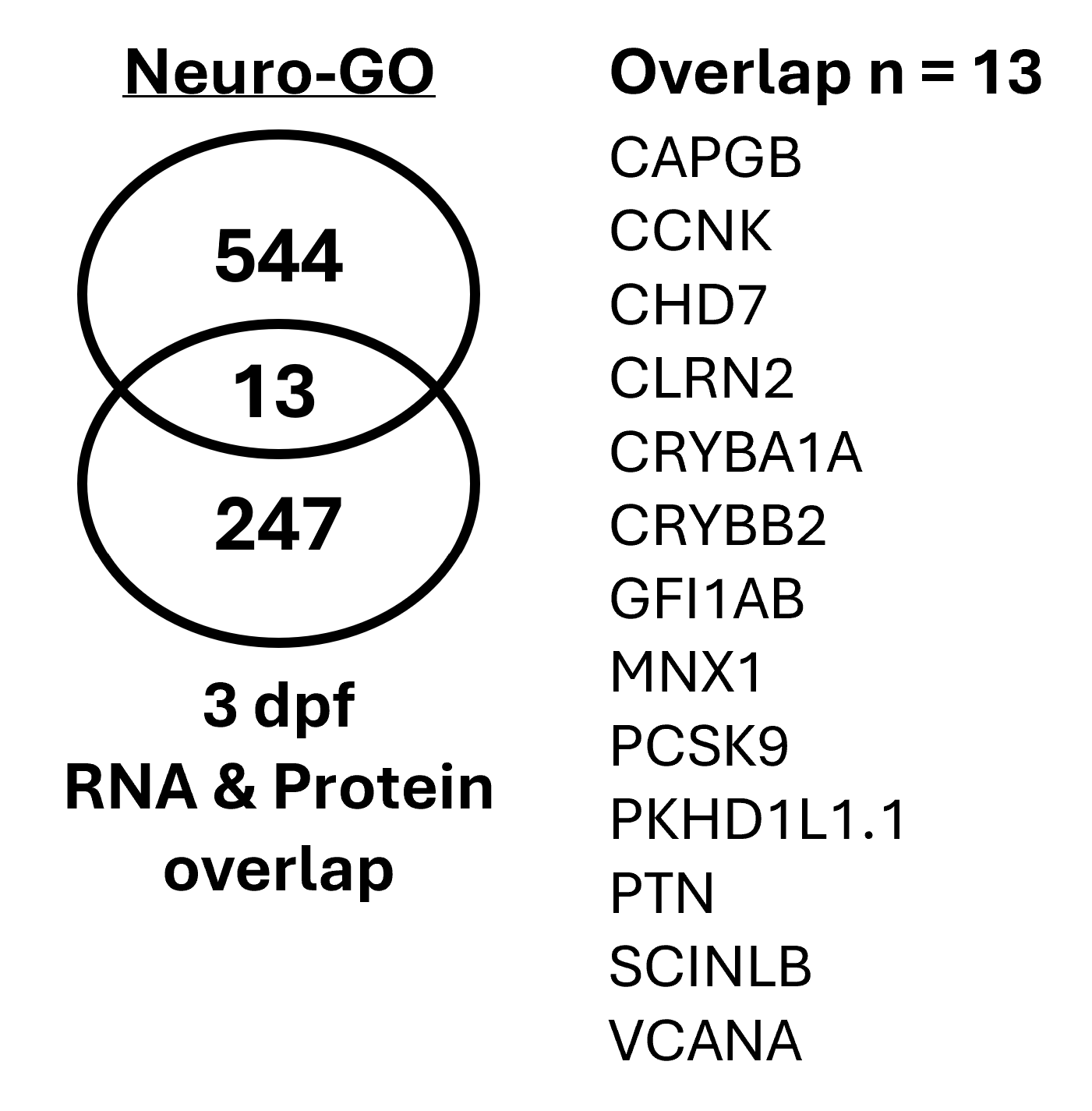

Supplemental Figure 6. List of gene symbol overlap between 5 dpf RNA HT v WT & MUT v WT and 5 dpf Protein HT v WT & MUT v WT from Figure 4K. List of gene symbol overlap between Neuro-GO list and 5 dpf RNA & Protein HT & MUT from Figure 4L.

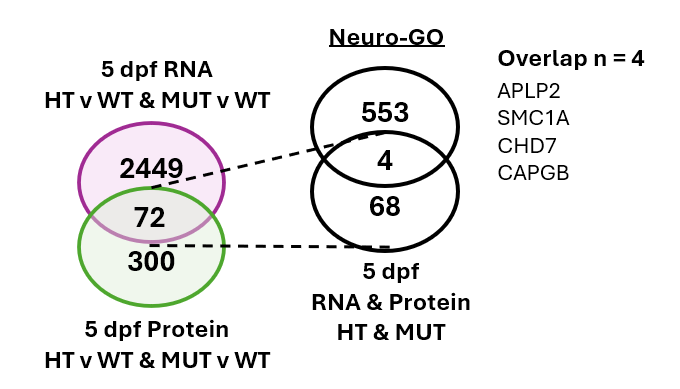

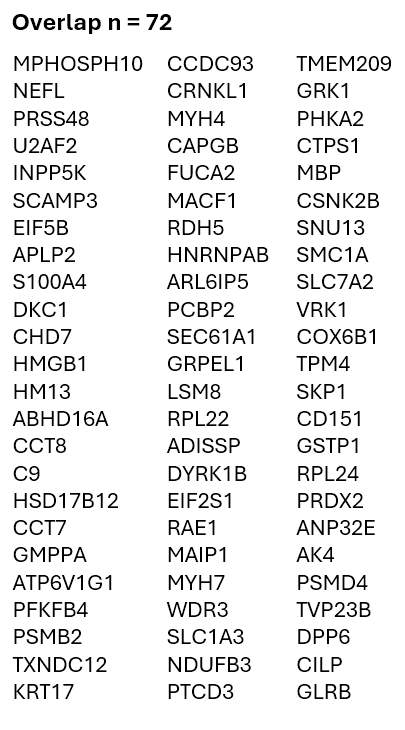

Supplemental Figure 7. List of gene symbol overlap between 3 dpf and 5 dpf, HT and MUT, RNA and Protein from Figure 5G.

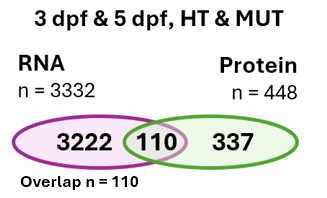

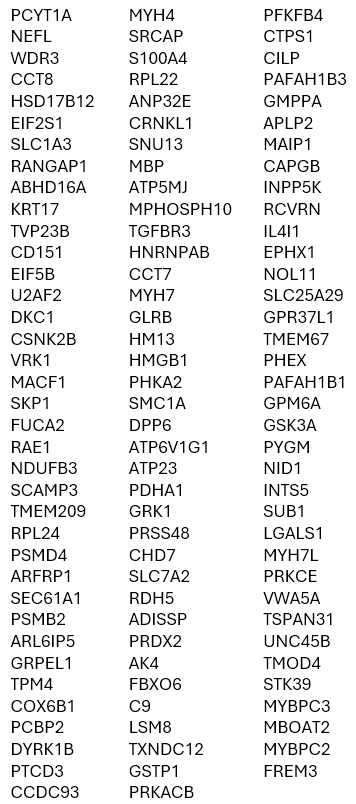

Supplemental Figure 8. List of gene symbol overlap between 3 dpf and 5 dpf, HT and MUT, RNA and Protein from Figure 5G with public ChIP-seq data from Schnetz 2010 from Figure 5H.

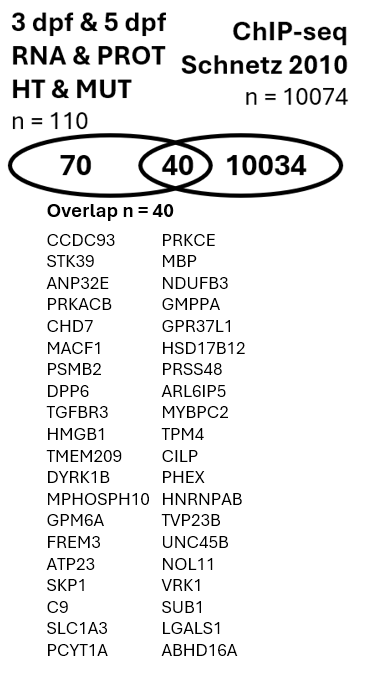

Supplemental Figure 9. List of gene symbol overlap between 3 dpf and 5 dpf, HT and MUT, RNA and Protein from Figure 5G with public ChIP-seq data from Reddy 2021 from Figure 5I.

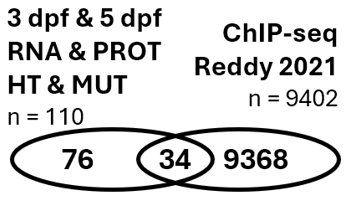

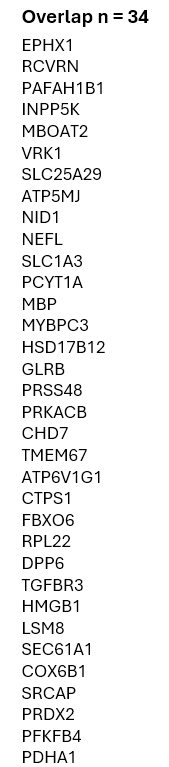

Supplemental Figure 10. Confirmation of CRISPR-Cas9 induced edits with designed gRNAs in candidate genes *capgb*, *nelfa*, *rdh5*, and *myh7* and no edits in candidate genes *lglals2b* and *pafah1b1a* by gel electrophoresis of candidate gene gRNA target sites (Table 1).

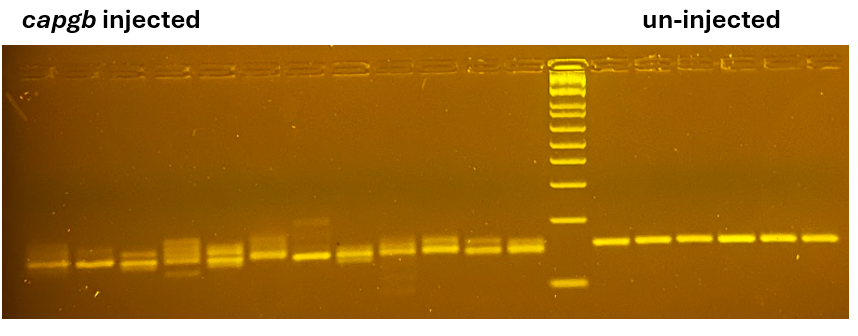

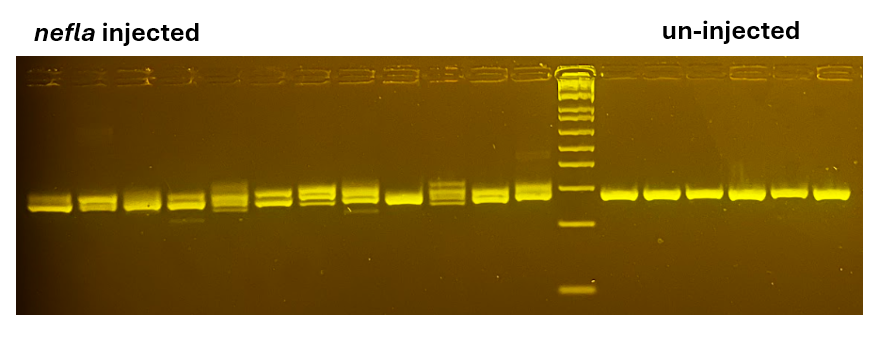

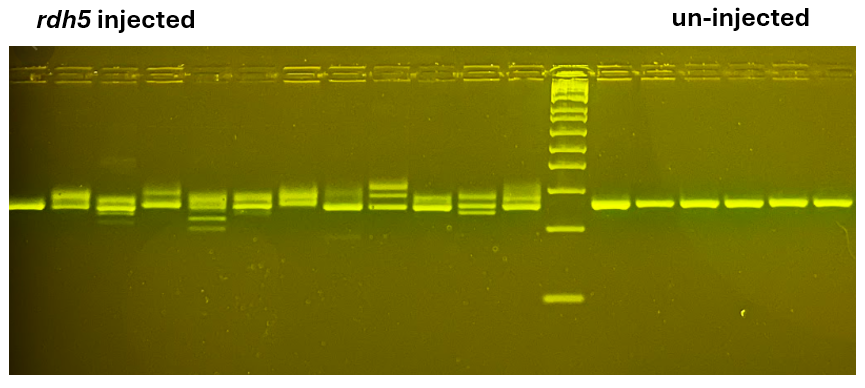

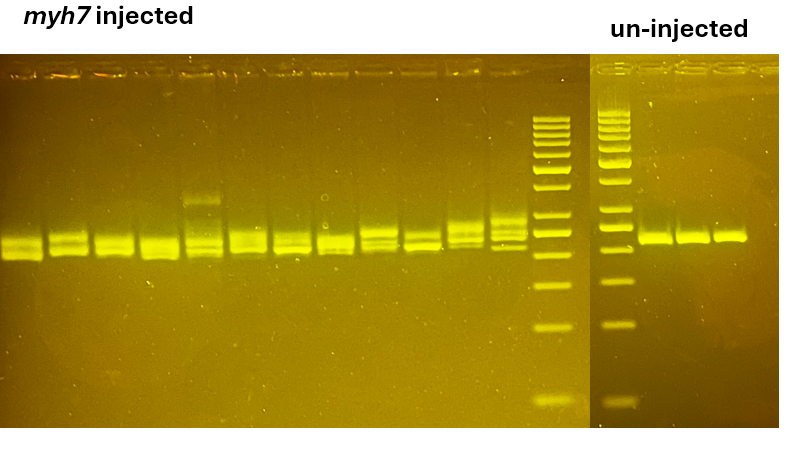

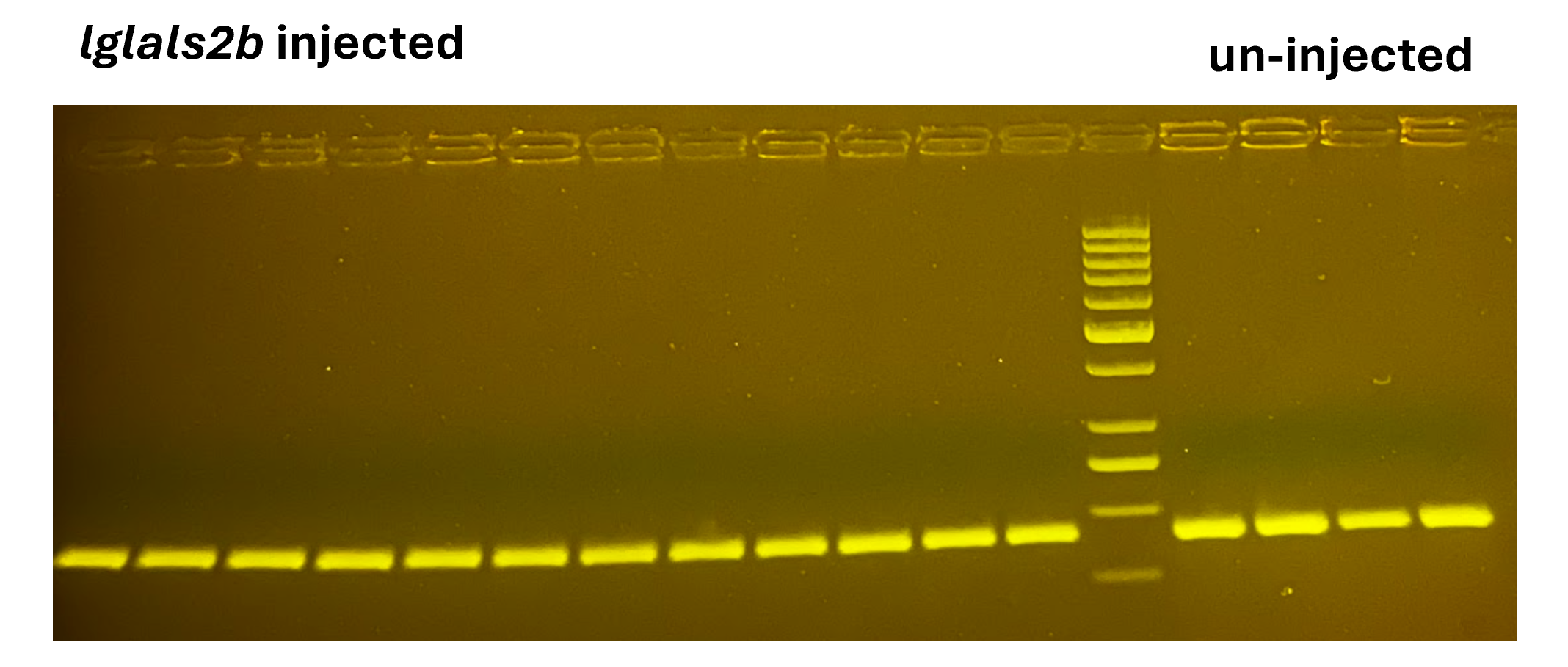

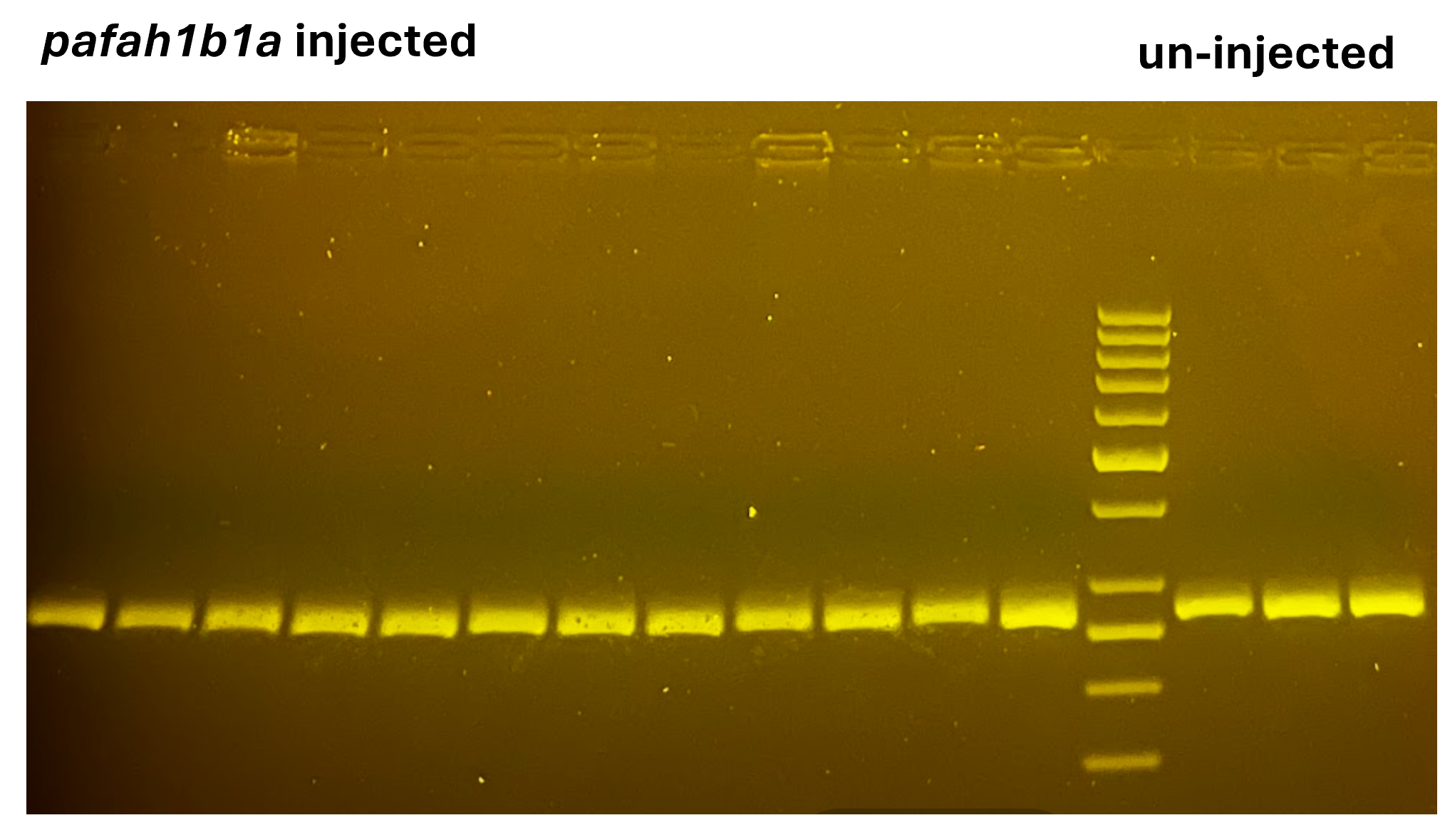

Supplemental Figure 11. Confirmation of no CRISPR-Cas9 induced edits with designed gRNAs in candidate genes *lglals2b* and *pafah1b1a* by sanger sequenced gRNA target sites (gray) aligned to Ensembl gene reference sequence, Snapgene version 7.2.1.

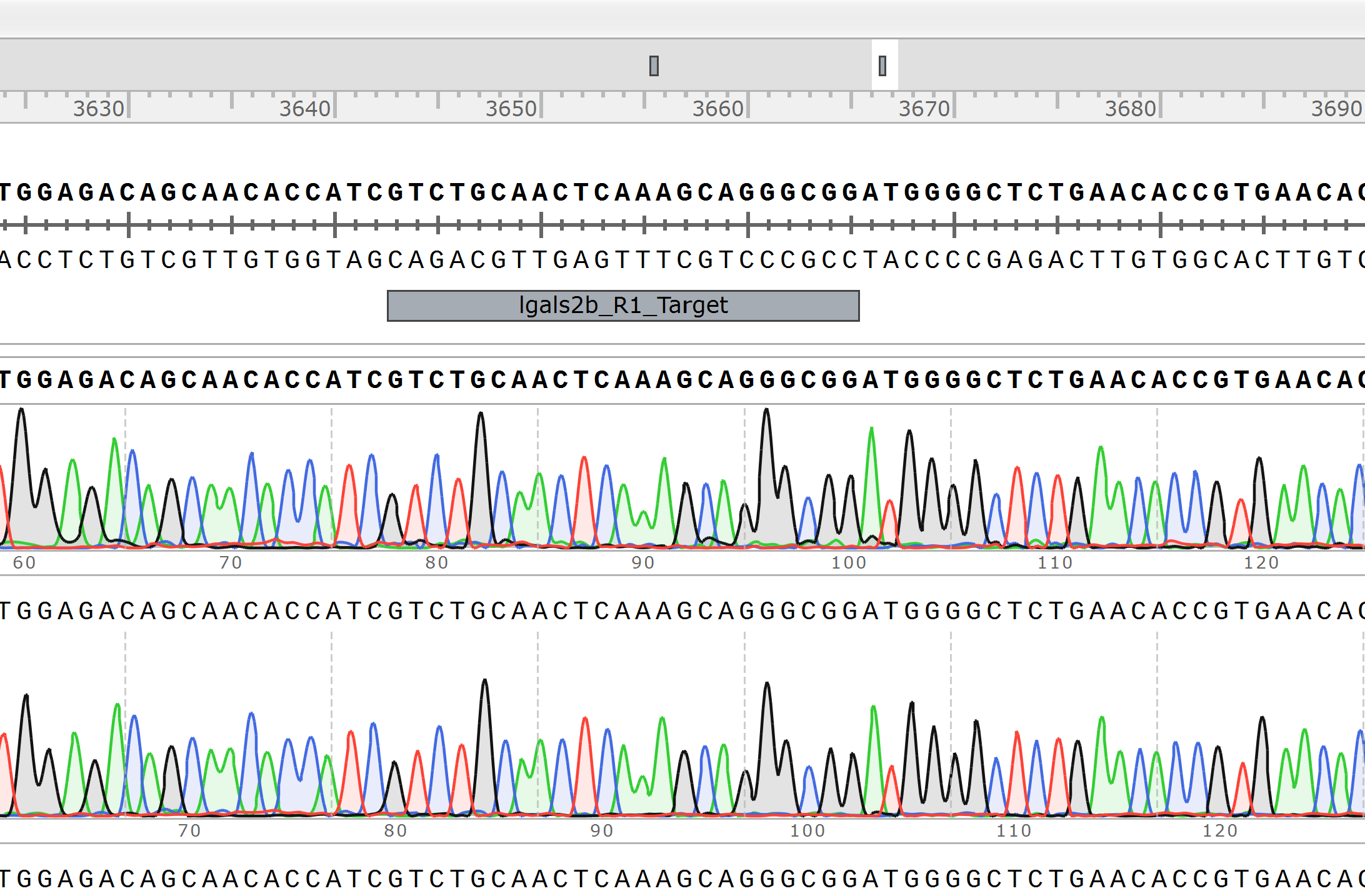

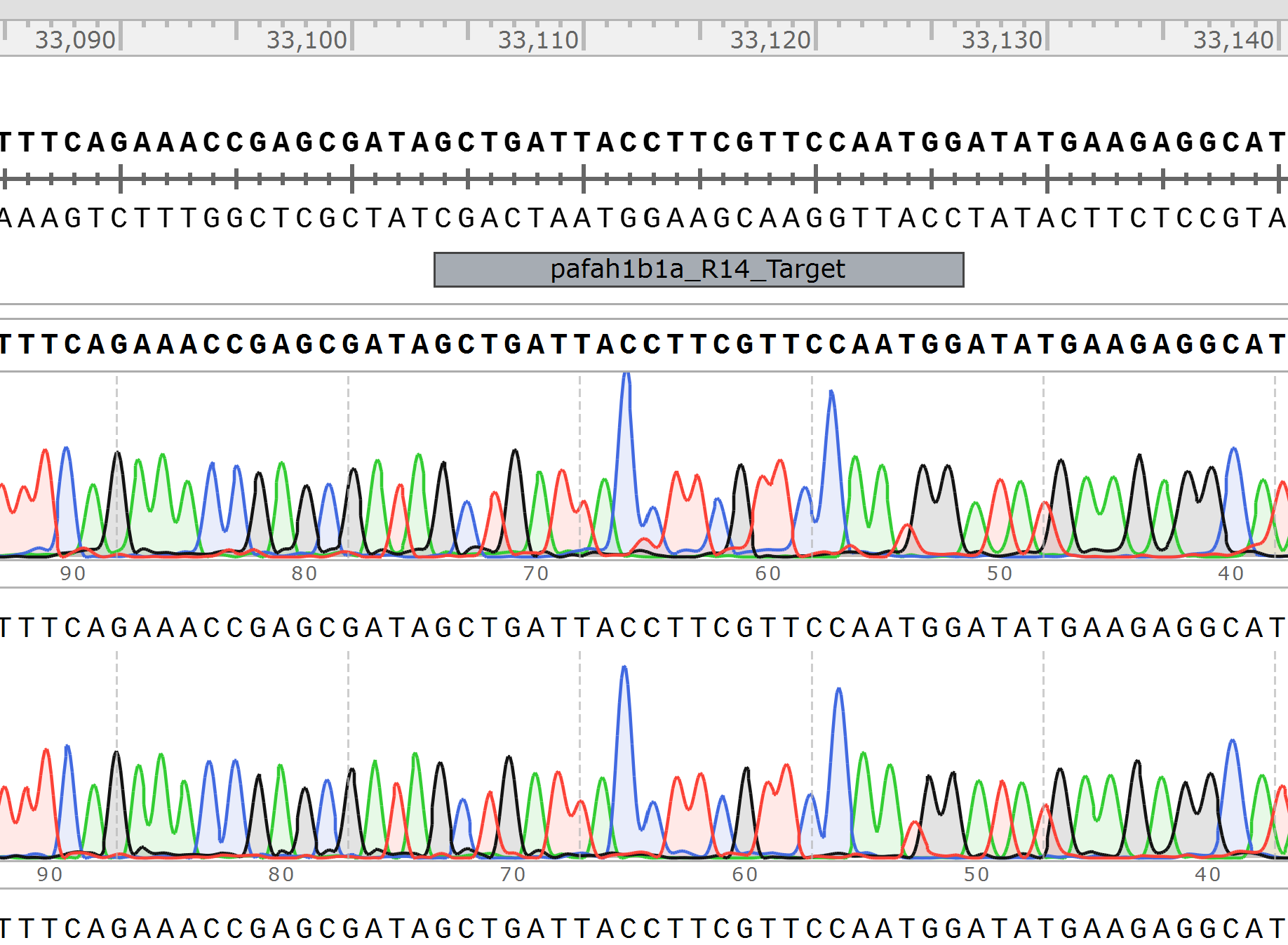

Supplemental Figure 12. Candidate gene symbols, Ensembl Gene IDs, number and percentage of CRISPR-Cas9 induced edits at least one gRNA target site.

| Gene Symbol | Ensembl Gene ID | Number of edits | Total | Percent edits |
| --- | --- | --- | --- | --- |
| *capgb* | ENSDARG00000099672 | 54 | 61 | 88.5% |
| *nefla* | ENSDARG00000057568 | 61 | 67 | 91.0% |
| *rdh5* | ENSDARG00000008306 | 67 | 70 | 95.7% |
| *myh7* | ENSDARG00000079564 | 23 | 23 | 100% |
| *lgals2b* | ENSDARG00000038153 | 0 | 64 | 0% |
| *pafah1b1a* | ENSDARG00000032013 | 0 | 33 | 0% |
